# Microsecond pulse electrical stimulation modulates cell migration

**DOI:** 10.1101/2022.10.23.513372

**Authors:** Xiao-Wei Xiang, Hao-Tian Liu, Wei Liu, Ze-Yao Yan, Yu-Lian Zeng, Ya-Jun Wang, Jing Liu, Yu-Chen Chen, Sai-Xi Yu, Cai-Hui Zhu, Xiao-Nan Tao, Chen Wang, Jin-Tao Wu, Yang Du, Xin-Xin Xu, Hai Gao, Yaming Jiu, Jiong Ma, Jian Qiu, Lingqian Chang, Guangyin Jing, Ke-Fu Liu, Yan-Jun Liu

## Abstract

Wound healing is a complicated process for maintaining skin integrity after injury, for which electrical stimulations (ES) are ascribed to promote wound healing by facilitating cell migration. Time-shortening of the stimulation treatment from current hours to minutes for efficient wound healing but free of cell damage in return, is however rather a challenge. Here, a novel mechanism of ultrashort pulse electric field (PEF), microsecond PEF at higher voltage, is proposed and realized to promote wound healing under a much short time (seconds) for the total treatment. We revealed that microsecond PEF regulated actin cytoskeleton reorganization and focal adhesion turnover, promoting fibroblasts migration in 2D cell cultures under the pulse stimulation. This accelerated fibroblast migration was accompanied by the mutual promotion with extracellular matrix (ECM) alignment in 3D microenvironments, which cooperatively benefit the eventual wound healing, and these findings were further confirmed by the enhanced skin wound healing in a classic mouse model. Additionally, we coined an actin- and collagen-dependent mechanism of microsecond PEF-mediated wound healing. The quantitative mechanism proposed here for our novel microsecond pulse electric filed (μsPEF) methodology orients the new practical electric treatment in a wide range of biomedical applications, such as wound healing, regenerative medicine, and tissue engineering.

## 1 Introduction

Wound healing is a complex physiological process consisting of multiple cellular events for maintaining skin integrity after injury^1^. Fibroblasts, as a critical cell type, migrating to the area of wounds, can reconstitute the various connective tissue components and restore the dermis to promote wound healing^2^. Hence, modulating the cell migration is a popular strategy to enhance the wound healing process^3^, for which applications based on exogenous growth factors (e.g., fibroblast growth factor, FGF) and bioactive materials (e.g., dermal substitute) are now used to rapidly increase the migratory capacity of fibroblasts towards efficient wound recovery^4,5^. However, tedious preservation and complicated production procedures in these bio-chemical treatments prompt further investigation of physical protocols as alternative therapeutics. As an endogenous electric current arises at the wound site, in situ electrical stimulation (ES) can provide a novel strategy to overcome the limitations of exogenous substances and promote the non-invasive healing of complex wounds^6,7^.

The stimulation intensity controlled by the electrical voltage is the first essential factor, considering efficiency for wound healing on one hand, and the practical clinical performance and tissue safety on the other hand. Researchers are currently making various attempts to stimulate or enhance wound healing by manipulating current forms, including continuous direct current (DC) and non-continuous DC^8,9^. Several studies have shown that increasing the DC electric field intensity from 50 to 200 V/cm can increase the migration speed of fibroblasts^10^, and further increasing the electric field intensity to 400 V/cm can reduce the stimulation time from hours to 1 h to achieve the same effectiveness^11^. Due to the tissue damage caused by continuous electrical stimulation, pulse electric field (PEF) is believed to be more promising by regulating currents of constant polarity flowing in an intermittent output with controlled pulse, which has been tested to be safer and more efficient^12^, and can suppress electrothermal, electro-physical and electro-chemical hazards that may produce cytotoxic effects^13,14^. Most clinical trials have demonstrated the excellent effects of PEF on wound healing^8,15^. This PEF stimulation has been proved to be better than DC-EF in increasing cell migration speed, promoting cell migration persistence^16,17^, and additionally PEF-treated fibroblasts expressed higher levels of type I collagen further benefiting the wound healing^18^. However, hours of exposure in the current stimulation inevitably affects cell viability due to an electrochemical reaction and the Joule heat effect in the medium ^19^. Whether the PEF stimulation could shorten the treatment time to minutes or even seconds without cell injury and promote long-time cell migration remains poorly understood. The mechanism may be different from electric field-dependent electrotaxis. It has been shown that cells can response rapidly to the external electric stimulation within microseconds and cell electroporation is one of these repsonse^20^. Therefore, inspired by the ultrafast electroporation mechanism, the answer may lie in transient electroporation dynamics of cells responding to the PEF stimulation. To prevent the harmful effects on cell viability, we modulated the voltage, ultra-short pulse width and seconds stimulation time for subsequent stimulation. And yet, there is no detailed evidence clarifying the relationship between short duration stimulation with resulting electroporation effect and wound healing.

In this study, we proposed a PEF simulation with higher voltage that shortened treatment time down to seconds level. Microsecond pulse width (μsPEF) was tailor-modulated in a novel way to prevent the cell damage with the preserved function of efficiently improving wound healing. To reveal the mechanism of how ultrashort pulse at high voltage promotes cell migration, μsPEF-induced fibroblasts migration in 2D and 3D cell cultures were quantitatively analyzed. We identified that 750 V/cm short-duration μsPEF instigated long-time and fast fibroblasts migration without cytotoxic effects. We also demonstrated that μsPEF stimulation regulated actin cytoskeleton reorganization and focal adhesion turnover to promote fast fibroblasts migration in 2D cell culture. Furthermore, we found that μsPEF could induce collagen fibers alignment in the 3D extracellular matrix, which provided fibroblasts contact guidance for morphological adjustment and fast cell migration, and this electrical stimulation could also promote the re-epithelialization process and remodeling phases of mouse skin tissues. Our novel μsPEF techniques could promisingly facilitate the wound healing and will find a wide range of biomedical applications in regenerative medicine and tissue engineering.

## 2 Results and Discussion

### 2.1 Enhanced cell migration and growth factor secretion under μsPEF

To study the μsPEF effect on cell migration during wound healing, we generated the rectangular pulsed monophasic μsPEF waveforms, by using self-built solid-state Marx generator (SSMG) to implement microsecond pulse with rising/falling edges at the nanosecond level for stimulation (Supplementary Fig. 1a). In the meanwhile, we built an electrical stimulation cell platform containing a customized six-well plate connected to this generator, compatible with various optical microscopes, to simultaneously track cell migration after μsPEF stimulation with different electric field intensities (Supplementary Fig. 1b). Institute for Medical Research-90 (IMR90), as a classical fibroblast cell line, was used to study the behaviors of fibroblasts during the procedure of wound healing under μsPEF (Fig. 1a).

**Fig. 1.**
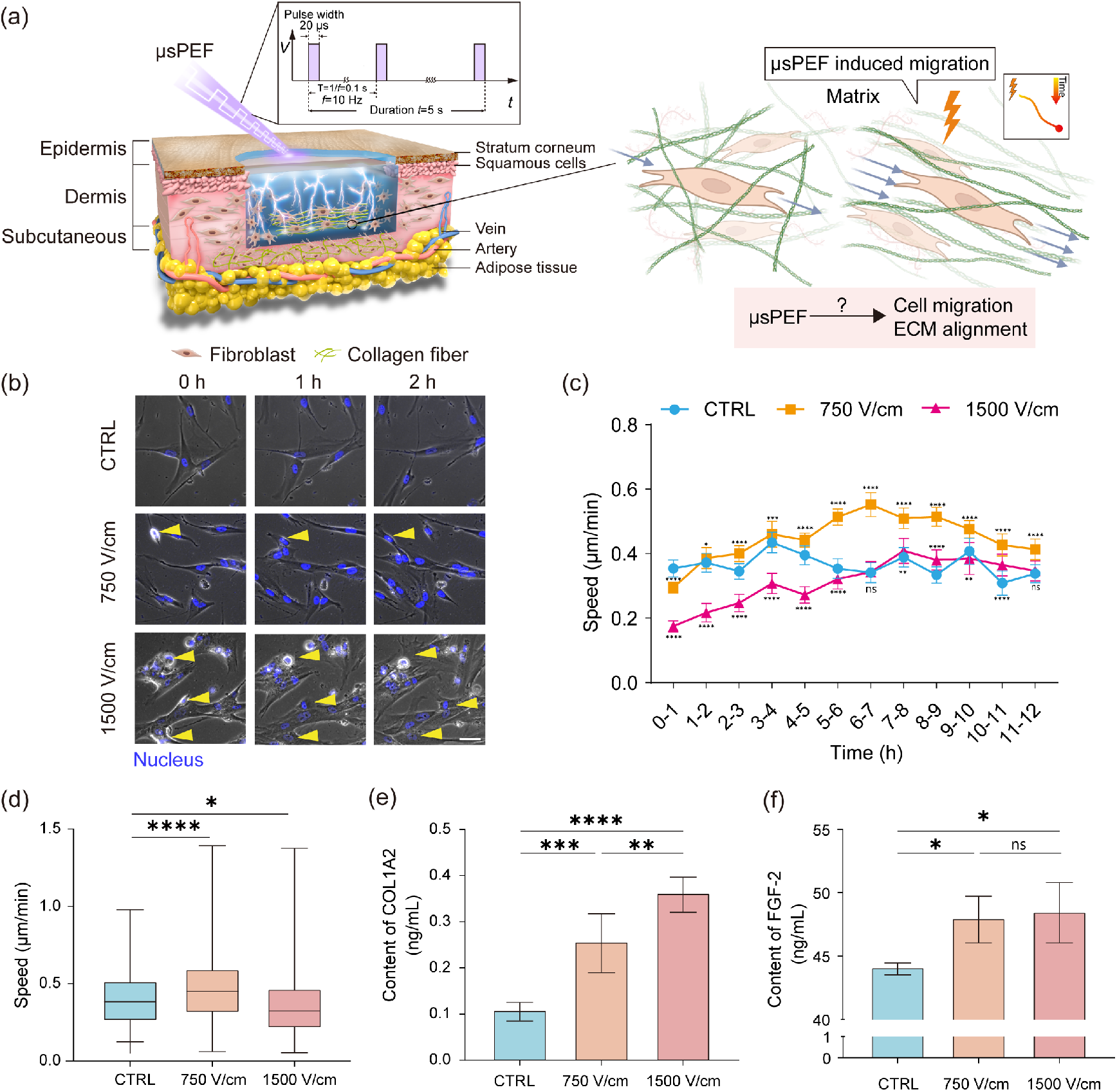
Enhanced cell migration and growth factor secretion under μsPEF. (a) A graphical illustration showed the effects of μsPEF (pulse width:20 μs, frequency: 10 Hz, duration: 5 s) on the skin wound, promoting cell migration and extracellular matrix remodeling. (b) Representative time-lapse images showing the fibroblasts morphology after μsPEF (i.e., 750 and 1500 V/cm) treatment of different intensity at different time points (i.e., 0, 1 and 2 h). Detached cells are marked by yellow arrow. Scale bar, 50 μm. (c) The line graph representing the cell migration average speed per 1 h from 0 to 12 h after different intensity μsPEF (i.e., 750 and 1500 V/cm) treatment (blue circle: control; orange square: 750 V/cm; pink triangle: 1500 V/cm). Results are presented as mean ± standard deviation with 95% CI (n_CTRL_=42 cells, n_750 v/cm_=48 cells, n_1500 v/cm_=47 cells); ^*^p<0.05, ^**^p<0.01, ^***^p<0.001, ^****^p<0.0001 versus control by one-way *ANOVA* for multiple comparisons. (d) Box plot showing the average cell migration speed with different intensity μsPEF (i.e., 750 and 1500 V/cm) and control treatments in 24 h. Results are presented as mean ± standard deviation with 95% CI (n_CTRL_=470 cells, n_750 V/cm_=979 cells, n_1500 V/cm_=535 cells). (e) Effects of μsPEF on the secretion of COLA2. The level of collagen type I α2 in cellular supernatants was measured after 48 h. The concentration of COLA2 was 0.109 ± 0.018 ng/mL in the control group, 0.257 ± 0.058 ng/mL in the 750 V/cm group and 0.363 ± 0.034 ng/mL in the 1500 V/cm group. Results are presented as mean ± standard deviation with 95% CI (n=5); ^*^p<0.05, ^**^p<0.01, ^***^p<0.001, ^****^p<0.0001 versus control by one-way *ANOVA* for multiple comparisons. (f) Effects of μsPEF on the expression of FGF2. The level of FGF2 in cellular supernatants was measured after 48 h. The concentration of FGF2 was 44.139 ± 0.360 ng/mL in the control group, 48.012 ± 1.488 ng/mL in the 750 V/cm group and 48.523 ± 1.944 ng/mL in the 1500 V/cm group. Results are presented as mean ± standard deviation with 95% CI (n=3); ns=0.7329, ^*^p<0.05, ^**^p<0.01, ^***^p<0.001, ^****^p<0.0001 versus control by one-way *ANOVA* for multiple comparisons.

To confirm that the short-duration electrical stimulation does not cause excessive Joule heating, we measured the temperature change on the medium under different electric filed strength stimulation (Supplementary Fig. 1c). As the electric field strength increases, the temperature rise increased within 5 ℃, where the electric field strength below 750 V/cm caused a temperature rise of no more than 1 ℃. Therefore, the temperature rise due to short-duration electrical stimulation can be ignored. In order to stimulate the cells to a greater degree in short time while maintaining their viability, we first optimized electrical parameters on fibroblast stimulation. In the meanwhile, we found that cell electropermeabilization occurred after μsPEF stimulation, which was verified by PI dyes transfer into cells (Supplementary Fig. 1d).

At 750 V/cm, fibroblasts maintained high perforation efficiency while having high cell viability, while at 1500 V/cm, the sum of non-viable and porated cells exceeded 100%, indicating that a fraction of electroporated cells did not restore membrane integrity. Therefore, 750 V/cm and 1500 V/cm were selected as typical values for inducing reversible and irreversible electroporation of fibroblasts, respectively. We then tested cytotoxicity of μsPEF on cells by measuring the NAD(P)H-dependent cellular oxidoreductase enzymes in the culture medium using MTT assay. The cell viability analysis demonstrated that there were no significant differences between the μsPEF exposed cells and the control at both day 1 and 3 after stimulation, meaning μsPEF with these two electric field strengths had no cytotoxicity effects on fibroblasts (Supplementary Fig. 1e). Next, we compared the cell morphology at different time points after μsPEF stimulation. We found that some cells detached immediately within 5 min after μsPEF stimulation, and the cells with 1500 V/cm stimulation expressed more severely detached than at 750 V/cm stimulation which were slowly returned to the adherent state over time (Fig. 1b). We further found that the adhesion ability of fibroblasts decreased with increasing μsPEF intensity, especially at 1500 V/cm stimulation and it remained stable after 1.5 h of electrical stimulation, which was consistent with the time point we observed for cell re-adhesion (Supplementary Fig. 1f). To explore whether the change in adhesion status affects the migration speed of the cells, we counted the speed each hour for 12 hours after μsPEF stimulation (Fig. 1c). The statistical analysis results showed that the migration speed of fibroblast cells significantly decreased within 1 hour after μsPEF stimulation. Then, the speed in the 750 V/cm group started to increase and surpassed the control after 2 hours stimulation which maintained at a high level for the following time. And the cell migration velocity upon the 1500 V/cm stimulation was steadily increasing, lower than control after 7 hours stimulation and it maintained a similar trend to control for the following time. It means that 750 V/cm stimulation could promote cell migration. We further tracked individual cell trajectories for 24 hours to investigate the influence of long-time migration behavior of fibroblasts after μsPEF stimulation (Supplementary Fig. 1g). Based on the analysis of the migration trajectory, fibroblast cells exposed to 750 V/cm stimulation presented a greater increase in cell migration speed while cells exposed to 1500 V/cm stimulation expressed a statistically remarkable decrease compared to control (Fig. 1d). Therefore, μsPEF could change cell morphology and thus affect cell migration behavior, with 750 V/cm μsPEF inducing faster migration than 1500 V/cm μsPEF. Zhao et al. represented that cell migration speed was significantly increased at a voltage of 200 or 400 mV/mm and a pulse width of 200 ms, compared to the DC EF-treated and control groups. In addition, PEF stimulation can maintain higher cell viability than DC EF^12^. Overall, a μsPEF stimulation could promote an increasement in the migration speed of cells over a long period of time greater than 12 hours, which has potential to significantly reduce clinical treatment time.

We then investigated the permeability of cell membrane at specific electric field strengths (Supplementary Fig. 1h). The fluorescence intensity increased logarithmically after μsPEF stimulation, and 1500 V/cm stimulation had the fastest intensity growing rate and highest thresholds among three groups, suggesting higher intensity of μsPEF stimulation generated the higher the cell permeability (Supplementary Fig. 1i). The scanning electron microscope (SEM) results showed that cell membrane underwent electroporation exposed to μsPEF. And cells treated with 1500 V/cm had larger membrane pores than those treated with 750 V/cm (Supplementary Fig. 1j, k). Thus, higher electric field intensities μsPEF stimulation on cells produced greater membrane perturbations, resulting in changes in cell permeability that lead to exchange of substances inside and outside the cell, which may promote enhanced extracellular protein secretion. Secretory proteins, type 1 collagen α2 (COL1A2) and fibroblast growth factor 2 (FGF-2), are particularly important in regulating cell migration behavior during wound healing^21,22^. Rouabhia et al. found that FGF-2 secreted by fibroblasts was significantly upregulated by 50 and 200 mV/mm ES more than 1 h long-term stimulation^10,23^. However, whether shorter stimulation could boost the secretion of extracellular protein remained unclear. To study whether μsPEF stimulation affect the fibroblasts secretion of COL1A2 or FGF-2, we detected their level in cellular supernatant after 48 hours stimulation by ELISA. The results showed that COL1A2 increased 137% and 233% after μsPEF stimulation at 750 V/cm and 1500 V/cm respectively, compared to control, thus promoting fibroblasts to synthesis new ECM to support other cells associated with effective wound healing (Fig. 1e). Meanwhile, the level of FGF-2 increased to 109% and 110% after μsPEF stimulation at 750 V/cm and 1500 V/cm respectively, acting as a survival factor in many models of cell and tissue injury by strongly activating fibroblast migration (Fig. 1f). Taken together, the results indicated that μsPEF stimulation not only affected the cell migration ability but also positively affected the secretion of collagen and growth factor.

### 2.2 μsPEF enhanced migration regulated by actin cytoskeleton rearrangement and focal adhesion turnover rate

Cell migration is a highly integrated process resulting from multiple, complex changes in intracellular organelles, mainly related to actin cytoskeleton and focal adhesions (FAs) ^24-26^. The applied external electric pulses demonstrate to be able to alter the actin cytoskeletal reorganization which affects the cell adhesion and migration^27,28^. To explore how μsPEF promoted cell migration, we analyzed the cytoskeleton and FAs dynamics in fibroblasts. The stability of the actin cytoskeleton in 24 hours after μsPEF treatment was confirmed by F-actin fluorescent images (Fig. 2a). As the μsPEF strength increased, the actin fibers became fragmented immediately (0 h), meanwhile its length and width decreased, and this decreasing effect was amplified with the increasing field intensity (Fig. 2b, c). Specifically, the cytoskeleton was only visible at the periphery of the cell stimulated with 1500 V/cm, and it showed a honeycomb-like appearance as previously observed. Several studies reported diminished F-actin features presumably indicating more actin depolymerization^29^. The actin cytoskeleton started to recover after 1 hour, the decreasing effects of μsPEF faded and the filaments were reconstructed completely under 750 V/cm stimulation, while filament with 1500 V/cm stimulation remained pale and fine after 24 hours (Fig. 2a). The cytoskeleton serves as a support for cells to maintain their morphology, and we found that cell area changed as the cytoskeleton expression, which indicated fibroblasts exposed to μsPEF leads to rapid but temporary disruption of actin filaments (Supplementary Fig. 2a, b). To study the dynamics of F-actin, we performed live-cell imaging of F-actin and found that the particle movement of F-actin increased significantly after 750 V/cm PEF treatment, while there was no significant change under 1500 V/cm stimulation compared to control (Supplementary Fig. 2c, d). Therefore, we concluded that μsPEF could induce actin cytoskeleton arrangement, which positively attribute to cell migration.

**Fig. 2.**
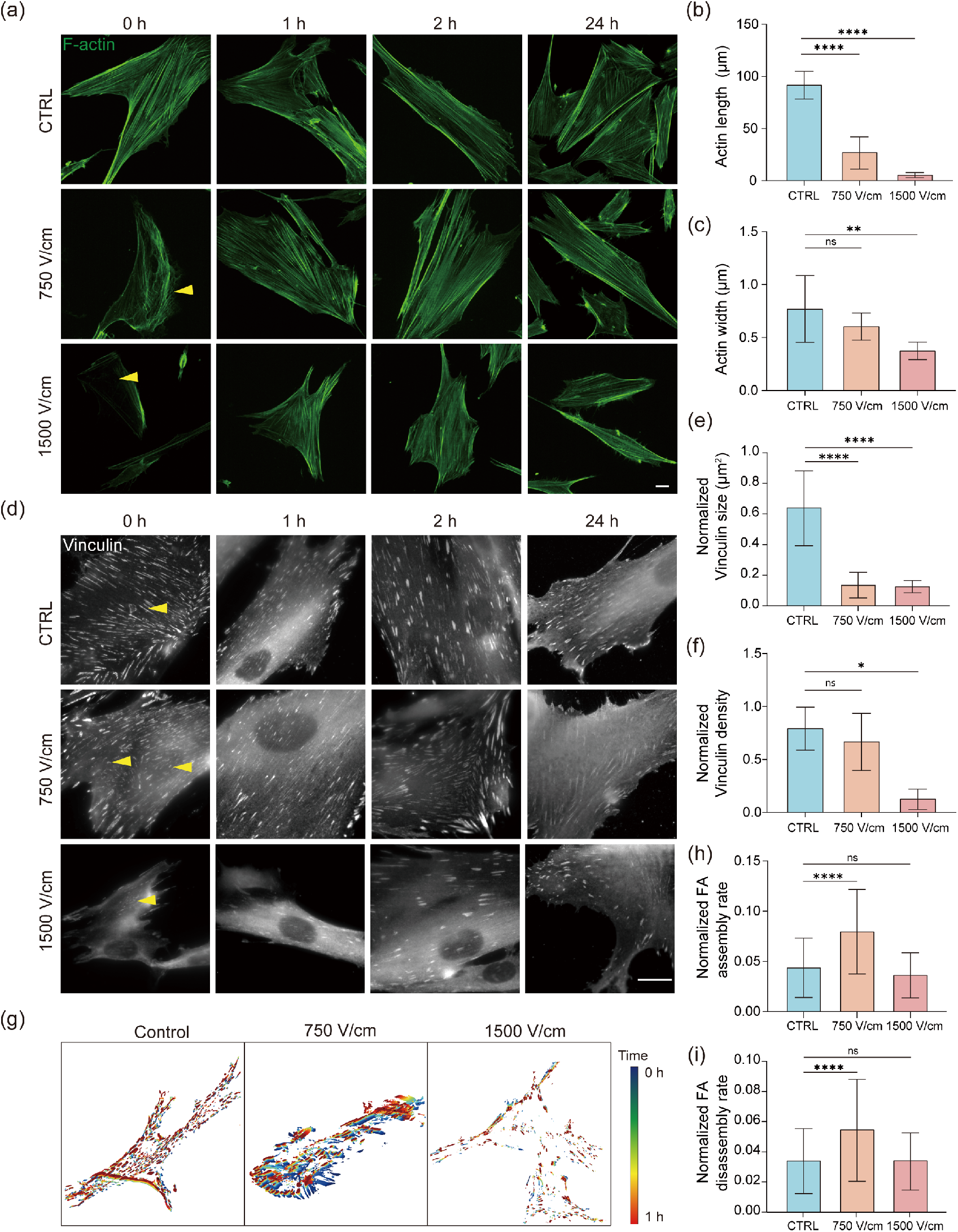
μsPEF enhanced migration regulated by actin cytoskeleton rearrangement and focal adhesion turnover rate. (a)Representative fluorescence images of IMR90 cells expressing F-actin after different intensity μsPEF (i.e., 750 and 1500 V/cm) treatment at different time points (i.e., 0, 1, 2 and 24 h). Cell membrane shows a honeycomb-like appearance (yellow arrow). Scale bar, 20 μm. (b)∼(c) Quantification of F-actin length and width upon μsPEF exposure (i.e., 750 and 1500 V/cm). Results are presented as mean ± SEM with 95% CI (n_CTRL_=7, n_750 V/cm_=9, n_1500 V/cm_=10). *p<0.05, **p<0.01, ns=0.2073 versus control by one-way ANOVA for multiple comparisons. (d) Representative fluorescence images of IMR90 cells expressing vinculin after different intensity μsPEF (i.e., 750 and 1500 V/cm) treatment at different time points (i.e., 0, 1, 2 and 24 h). Focal adhesions show immature appearance (yellow arrow). Scale bar, 20 μm. (e)∼(f) Quantification of vinculin area and density upon μsPEF exposure (i.e., 750 and 1500 V/cm). Results are presented as mean ± SEM with 95% CI (n_CTRL_=3 cells, n_750 V/cm_=3 cells, n_1500 V/cm_=3 cells). ^*^p<0.05, ^****^p<0.0001, ns=0.7265 versus control by one-way *ANOVA* for multiple comparisons. (g) Representative images of FA longevity analyzed by FAAS, with each focal adhesion outlined in a color according to the occurring time in the supplementary movie 3. The adhesions in blue have the longest lifetime, whereas those in red have the shortest longevity. (h) ∼(i) Bar charts showing normalized FA assembly and disassembly rate of cells in the control and μsPEF group. Results are presented as mean ± SEM with 95% CI (n=5 cells). ^***^p<0.001, ^****^p<0.0001, ns_assembly_=0.3931, ns_disassembly_>0.9999 versus control by one-way *ANOVA* for multiple comparisons.

FAs act as mechanosensors that can also reflect the cellular response to μsPEF stimulation at these sites in association with the actin cytoskeleton (Supplementary Fig. 2e). To investigate the dynamics of FAs, we imaged the FAs protein vinculin by Total Internal Reflection Fluorescence (TIRF) microscope, and statistically analyzed their changes in the size and density at different time points after stimulation (Fig. 2d-f). We noticed that the size of individual vinculin becoming smaller with increasing μsPEF intensity, which implied that more immature FAs being generated would affect cell adhesion. Interestingly, at the intensity of μsPEF of 1500 V/cm, the density of FAs proceeded a significant decrease, while at 750 V/cm, the density of FA remained unchanged from the control (Fig. 2e, f). This phenomenon corresponded to the cell detachment that cell rounding occur to accommodate changes due to breakdown of adhesion sites, and the results further validated that at higher electrical stimulation was able to lead to decreased cell adhesion (Fig. 1b). The size and number of vinculin gradually recovered over a period of time after μsPEF stimulation, indicated that FAs were maturing. To further demonstrate the FAs dynamics, we analyzed FAs assembly and disassembly rates from FAs turnover rate of living-cell imaging and FA Analysis Server (FAAS), respectively (Fig. 2g). The results showed that the rate of FAs assembly and disassembly was significantly increased under 750 V/cm μsPEF stimulation, however, there was no significant difference between the stimulation at 1500 V/cm and control (Fig. 2h, i). Therefore, the 750 V/cm treated cells had a high FA turnover rate. Based on the phenomena observed so far, we can speculate that μsPEF promoted cell migration mainly by enabling the reorganization of the actin cytoskeleton and accelerating the turnover rate of FAs.

### 2.3 Collective cell directional migration in response to μsPEF

The tissue regeneration process involves collective cell migration at the wound edge. To investigate the effect of μsPEF on collective cells, we utilized a wound healing assay to study collective cell migration (Fig. 3a). These wound area time-lapse images and quantification analysis showed the fastest decrease in wound area under 750 V/cm stimulation, and relatively slow migration rates with 1500 V/cm stimulation and control (Fig. 3b, c). To fully describe the displacement of the collective cells, we used particle image velocimetry (PIV) to express the displacement fields which showed migration vectors’ speed and direction, and these vectors were not always towards the free surface, where the collective cells under 750 V/cm stimulation tended to migrate more toward the free surface (Fig. 3d). In order to better understand the distribution of the activity vectors in the representation space, we applied t-SNE to map the representation vectors in the dataset to 2D and 3D visualizations (Supplementary Fig. 3a, b). The 2D visualization clearly showed two clusters after adding the vector direction factor, and no overlap was observed in the 3D visualization, which indicated that collective migration could be divided into forward migration (0°≤ flow direction angle <180°) and backward migration (180°≤ flow direction angle <360°). To further analyze the directionality of collective migration, we considered collective migration as a migration flow and obtained the frequency distribution according to the flow direction (Supplementary Fig. 3c). The statistics results indicated that the frequency of forward movement toward the free surface was more significant than the backward movement. Moreover, we extracted the speed of the forward movement to discriminate the effect of μsPEF stimulation on directional collective migration. We found that the migration speed was significantly higher after 750 V/cm stimulation, while the speed was decreased after 1500 V/cm stimulation compared to control (Supplementary Fig. 3d). Therefore, these results demonstrated that 750 V/cm μsPEF stimulation resulted in faster collective cells migration to the free surface, thereby promoting wound closure.

**Fig. 3.**
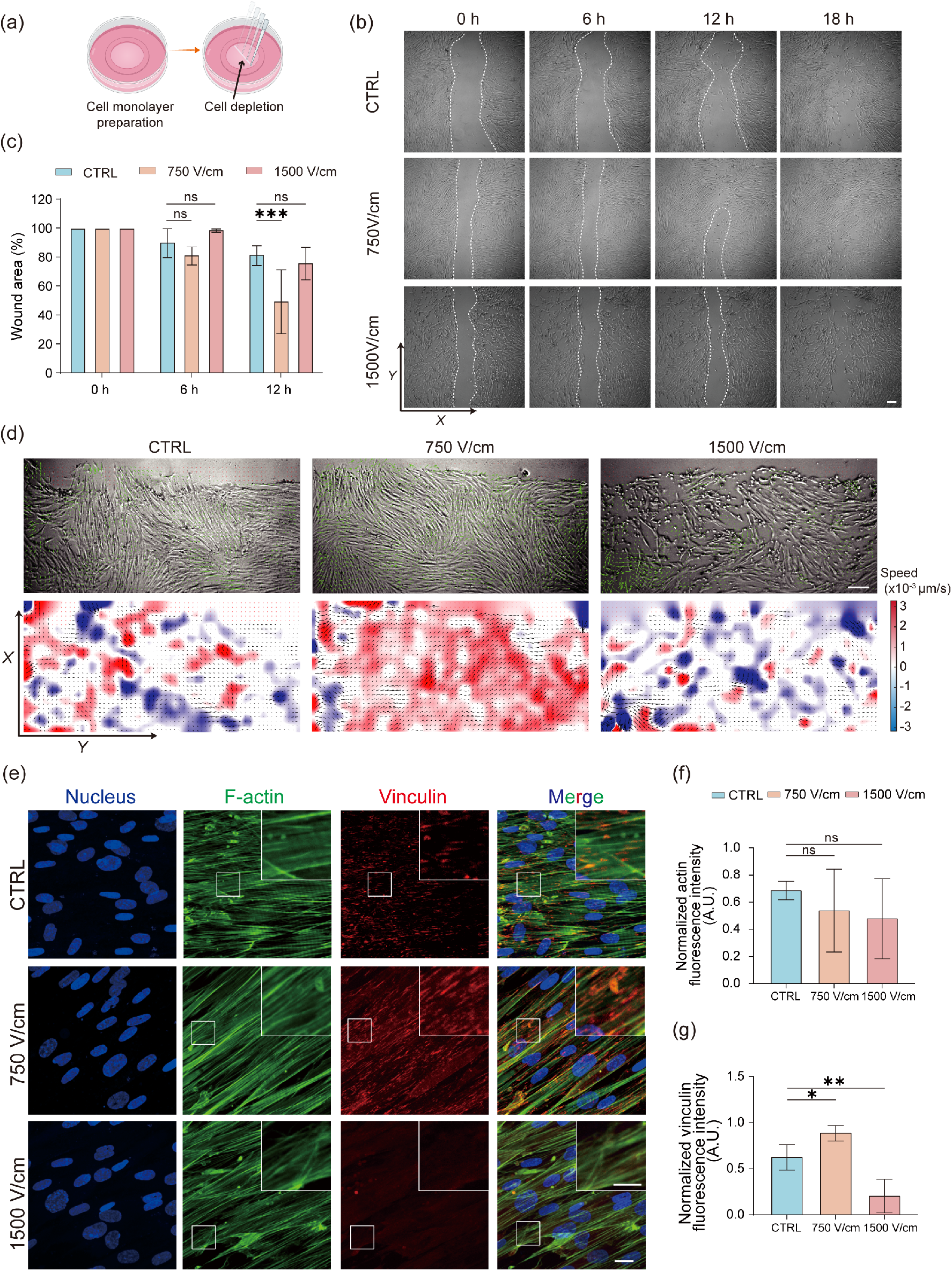
Collective cell directional migration in response to μsPEF. (a) A graphical illustration showing the process of the wound healing assay. Cells were scratched away from the monolayer using a tip. Created with BioRender.com. (b) Time-lapse microscopic images comparing IMR90 wound healing assays with and without μsPEF stimulation, starting at 0 h and lasting for 24 h after stimulation. Scale bar, 200 μm. (c) Graph showing the wound area covered by IMR90 over time. Under electrical stimulation, 750 V/cm enhanced wound healing at the fastest rate. Results are presented as mean ± SEM with 95% CI (n=3 experiments). ^*^p<0.05, ns=0.9730 versus control by unpaired Student’s *t* test. (d) Cell migration of monolayer fibroblast cells after different intensity μsPEF exposure (i.e., 750 and 1500 V/cm). It expands towards the empty space (direction of the increasing *X*). Top panel shows phase-contrast images of fibroblasts, taken at t=30 min. Corresponding 2D fields of cell velocity are shown in the bottom panel. Scale bar, 200 μm. (e) Representative fluorescence images of monolayer IMR90 cells with or without electrical stimulation. Scale bars, 20 μm and 10 μm (zoom). (f) Quantification of fluorescence intensity of F-actin after μsPEF exposure (i.e., 750 and 1500 V/cm). Results are expressed as mean ± SEM with 95% CI (n=3). ns_actin_750 V/cm_=0.6408, ns_actin_1500 V/cm_=0.4003 versus control by one-way *ANOVA* for multiple comparisons. (g) Quantification of fluorescence intensity of vinculin after μsPEF exposure (i.e., 750 and 1500 V/cm). Results are expressed as mean ± SEM with 95% CI (n=3). ^*^p<0.05, ^**^p<0.01 versus control by one-way *ANOVA* for multiple comparisons.

The most distinctive feature of monolayers compared to single cells is the cellular junctions, organized as a network of multicellular strands^30^. To analogize the difference between the electric field applied to single cell and collective cells, we built the electrical equivalent circuit (EEC) model for single and collective cells based on the previous description^31^ with *R*_*e*_, *R*_*i*_ and *C*_*m*_, as the resistance and parallel capacitance of the extracellular fluid, cell membrane, and intracellular fluid, respectively (Supplementary Fig. 3e). The simulation results showed that electric field distortion can be seen at the intercellular contact point of collective cells (Fig. 3b, Supplementary Fig. 3f). However, there was relatively uniform distribution of electric potential within single cells and collective cells (Supplementary Fig. 3f, g). Therefore, we could draw an analogy between the case of single cells and collective cells. Next, we found that there was no significant difference in the collective cells F-actin expression under these strengths, while vinculin expression significantly decreased with 1500 V/cm stimulation compared to control (Fig. 3e-g). This trend was consistent with the results of previous experiments, thus the response of single cell and collective cell after μsPEF stimulation were analogized. Overall, we found similarities in the electric field distribution between single cells and collective cells through the simulation results of the EEC model, and through the quantitative results of vinculin expression, we also found similar trends between single and collective cells after μsPEF stimulation, with reduced cell adhesion at 1500 V/cm stimulation, which would inhibit collective cell migration. And in terms of migration behavior, similar to the previous results for single cells, 750 V/cm μsPEF could promote collective cell migration, and additionally we found some evidence of directional migration, where 750 V/cm μsPEF can promote collective cell orientation to the free surface (forward movement).

### 2.4 μsPEF induced ECM remodeling to promote cell migration

Cell microenvironment in a living tissue is highly complex, involving cell-matrix interaction and cell-cell interaction. The components of ECM are key elements of the 3D microenvironments and are actively involved in the healing process by creating a scaffold that provides the structural integrity of the matrix^32,33^. To better mimic the physiology of the original tissue, we chose type I collagen, the most abundant protein of ECM components, to reconstitute 3D cell microenvironments for studying cell migration after μsPEF treatment. First, we investigated whether μsPEF affects the ECM, in term of its fiber alignment and matrix stiffness. The second-harmonic generation (SHG) microscopy images showed that collagen fibers underwent significant fiber reorientation or alignment parallel to the direction of the electric field lines after exposure to μsPEF (Fig. 4a). It has been reported that electric field could alter the biomaterial ultrastructure, allowing collagen fibril alignment; on the other hand, these aligned fibers have contact guidance on cell to direct migration^34,35^. To further investigate the collagen fiber feature, we extracted the length and angle of individual collagen fibers by CT-FIRE algorithm (Fig. 4a, b, Supplementary Fig. 4a-d). These quantitative results indicated that the alignment angle of collagen fibers tended to be homogenized after μsPEF stimulation, and this feature would produce contact guidance for cells and thus promote cell migration (Fig. 4b). We also found that the length, width and straightness of fibers suffered a decrease in response to μsPEF stimulation, and thus fiber degradation may have occurred (Supplementary Fig. 4b-d). Next, to investigate whether μsPEF stimulation degraded collagen and thus caused changes in these features, we quantified the concentration of collagen changes after electrical stimulation, the results of SDS-PAGE showed that the final concentration of collagen gel with the same initial concentration decreased with the increase of electric field intensity μsPEF stimulation, which meant that collagen gel underwent different degrees of degradation in response to different intensity μsPEF stimulation (Supplementary Fig. 4e, f). Then, we measured the stiffness of collagen after μsPEF stimulation by atomic-force microscopy (AFM), and the Young’s modulus increased under 750 V/cm stimulation compared with control (Fig. 4c). It has been shown that pulsed electrical signals can locally accelerate the nucleation of calcium phosphate nanocrystals in or on collagen and drive collagen calcium phosphate mineralization^36^. In addition, cells could mechano-sense matrix stiffness as they could apply contractile forces to fibrillar collagen, and there were reported that cells have durotaxis properties on stiffness gradients surface and migrate faster on harder surfaces^37^. Therefore, collagen fiber alignment and stiffness both increased by μsPEF stimulation will provide the support and guidance for 3-dimensional (3D) cell migration.

**Fig. 4.**
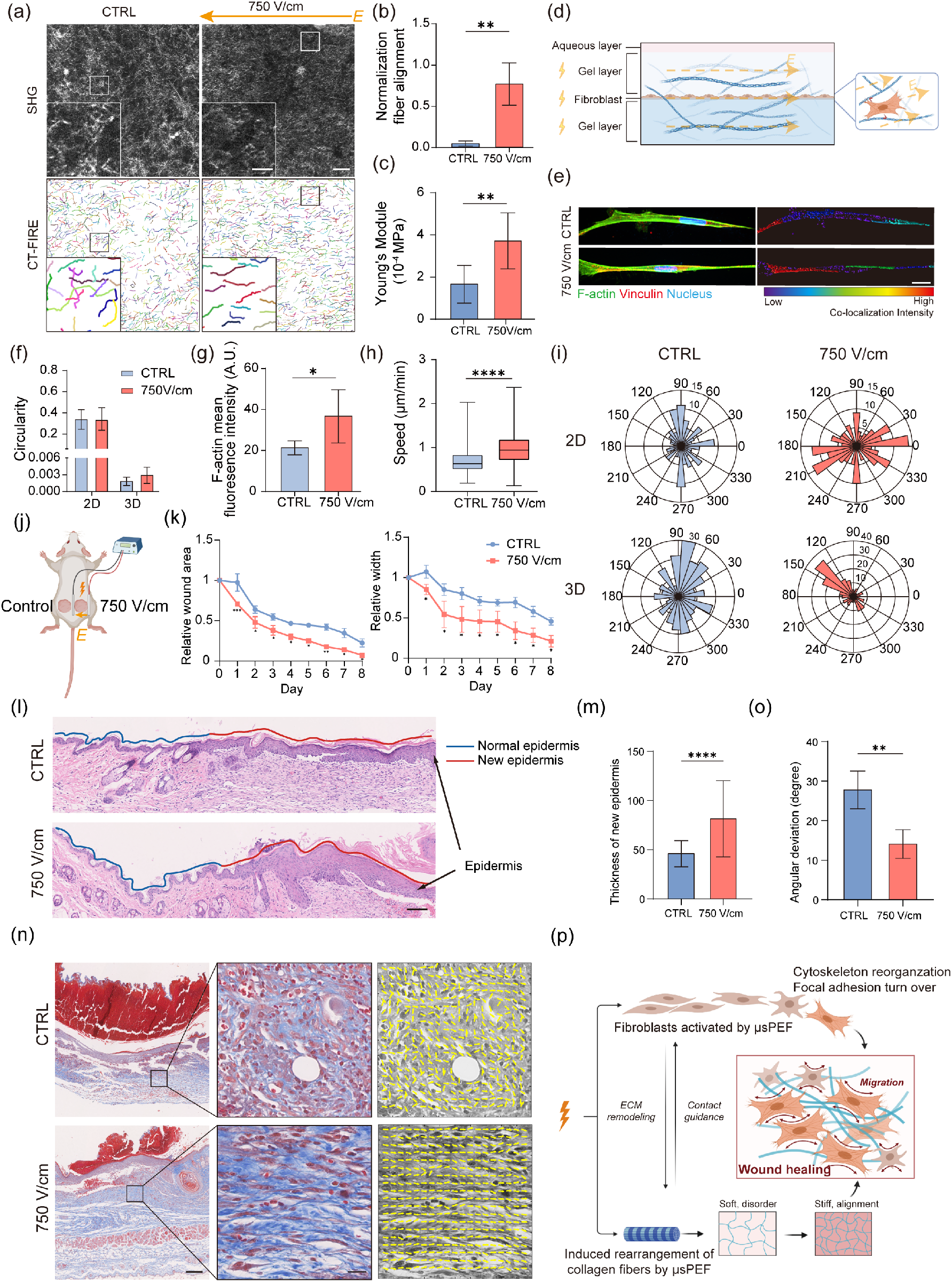
μsPEF induced ECM remodeling to promote cell migration. (a) Collagen fiber distribution and orientation in the control group and 750 V/cm μsPEF group. Top: Representative label-free second harmonic generation microscopy images of collagen fibers. Scale bars, 50 μm and 15 μm (zoom). Bottom: Representative images of collage fiber analysis using the open-source software CT-FIRE. The orange arrow is the direction of the electric field lines. (b) Bar chart showing quantitative analysis of collagen fiber alignment. Results are presented as mean ± SEM with 95% CI (n=3). ^**^p<0.01, versus control by unpaired Student’s *t* test. (c) Bar chart showing Young’s Modulus of IMR90 with and without μsPEF. Results are presented as mean ± SEM with 95% CI (n_CTRL_=8, n_750 V/cm_=11). ^**^p<0.01, versus control by unpaired Student’s *t* test. (d) A graphical illustration showing three-dimensional (3D) “collagen sandwiches” for IMR90 cells in 3D collagen gels under the stimulation of μsPEF. Created with BioRender.com. (e) Immunofluorescence analysis showing the expression of F-actin and vinculin in 3D-cultured IMR90 fibroblasts with or without electrical stimulation. Left: Representative fluorescence images of IMR90 in 3D collagen matrix. Right: Quantification of the colocalization of F-actin and vinculin by IMARIS. Color spectrum indicates the intensity of colocalization, with areas of low intensity in blue and high intensity in red. Scale bar, 30 μm. (f) Bar chart showing quantification of IMR90 cell circularity under 750 V/cm μsPEF exposure or control condition in the 2D and 3D environment. Results are presented as mean ± SEM with 95% CI (n=10 cells). (g) Bar chart showing the quantification of fluorescence intensity of F-actin in 3D-cultured IMR90 under different treatment. Results are presented as mean ± SEM with 95% CI (n=3). ^*^p<0.05 versus control by unpaired Student’s *t* test. (h)∼(i) Quantitative analysis of migration parameters for IMR90 in 3D collagen matrix under 750 V/cm μsPEF and control, including mean migration velocity and migration direction (compared with 2D). Results are presented as mean ± SEM with 95% CI (For mean migration velocity n_CTRL_=1101 cells, n_750 V/cm_=240 cells; for migration direction, n_2D_CTRL_=94 cells, n_2D_750 V/cm_=196 cells, n_3D_CTRL_=220 cells, n_3D_750 V/cm_=200 cells); ^****^p<0.0001 versus control by unpaired Student’s *t* test. (j) A graphical illustration showing full-thickness skin defect mouse model. In this model, two full thickness wounds are created on both sides of the midline allowing each mouse to serve as their own control. Left: control group, right: μsPEF treatment group. (k) Dynamic changes of wound area and width. Lines and error bars indicate mean and SEM. N=7, ^*^p<0.05, ^**^p<0.01, ^***^p<0.001 versus control by one-way *ANOVA* for multiple comparisons. (l) H&E-staining of wounded skin sections in different groups at day 8 post-operatively. Blue line: normal epidermis; Red line: new-born epidermis. Scale bar, 100 μm. (m) Quantitative analysis of the thicknesses of new-born epidermis layer using ImageJ. Results are presented as mean ± standard deviation; n = 7 for each group. ^****^p<0.0001 versus control by unpaired Student’s *t* test. (n) Masson staining of wounded skin sections in different groups at day 8 postoperatively. Left: Representative Masson images of full-thickness wounds. Middle: local zoom of the wounds. Right: Representative images of fiber orientations analyzed by the open-source software MatFiber. Scale bars, 200 μm and 20 μm (zoom). (o) Bar chart showing quantitative analysis of fiber angular deviation. Results are presented as mean ± SEM with 95% CI. ^**^p<0.01 versus control by unpaired Student’s *t* test. (p) Schematic summary of the main findings showing the effects of μsPEF on IMR90 migration and cellular mechanics. IMR90 cell activation under μsPEF is reflected in the accelerated rate of cytoskeletal movement as well as the turnover rate of vinculin, which promotes cell migration. The extracellular matrix undergoes rearrangement in response to electrical stimulation, which provides contact guidance and stiffness support to cells, thereby better facilitating cell migration in a 3D environment and *in vivo*. Created with BioRender.com.

Second, we investigated the migration behavior of cells in the ECM after μsPEF stimulation, we seeded fibroblasts in the 3D sandwich collagen scaffold to obtain better live cell observation in the same focal plane (Fig. 4d). The fluorescence images and circularity statistical results showed that cells exhibited more spindle like shape in the 3D environment compared to 2D (Fig. 4e, f). To explore the distribution of cytoskeleton (F-actin) and FAs (vinculin) of cells in a 3D environment after μsPEF stimulation, we performed co-localization images analysis between F-actin and vinculin within cells by IMARIS and found that vinculin would accumulate at the front of the cell to provide the support for the cell. Then we also found the increased expression of F-actin in the 3D environment when cells were stimulated at 750 V/cm μsPEF, which would provide cell more support thus promoting cell migration (Fig. 4g). To further analyze changes in cell migration behavior after μsPEF stimulation, we tracked individual cell trajectories for 24 hours, and found that the speed of 3D cell migration remarkably increased after 750 V/cm stimulation (Fig. 4h). In the meanwhile, the convergence of cell migration direction was found in μsPEF-exposed ECM, indicating that μsPEF induced cell directional migration in 3D, while cells in the unstimulated 3D and 2D environment still maintained the persistent random walk (PRW) movement pattern (Fig. 4i). Thrivikraman et al. found that aligned and cross-linked fibrin gel has an enhanced contact guiding effect on fibroblasts^38^. Also by micropatterning parallel lines of fibronectin to mimic fibrillar geometry, researchers found that fibroblasts elongated and preferentially aligned with the ECM, and the cell trajectories also became oriented parallel to the ECM^39^. Therefore, we concluded that ECM fiber alignment generated by μsPEF stimulation provided more guidance on the directionality of cell migration, thereby facilitating faster cell migration.

To investigate the potential of fibroblasts for ECM remodeling, we utilized the gel deformation assay by analyzing the volume changes of dissociated collagen gels that were embedded with cells to probe cell contractility. Interestingly, the time-lapse images and area analysis showed that cells were able to deform the gel faster with 750 V/cm stimulation compared to control. It demonstrated that the enhanced contractile force of the cell in response to 750 V/cm μsPEF (Supplementary Fig. 4g, h). When the cells in collagen exert contractile force, the cells will enrich the surrounding fiber more around the cells. The results of fiber and cell co-localization images showed that intense and oriented collagen fiber bundles were particularly noticeable around the cell periphery (Supplementary Fig. 4i). The degree of co-localization after μsPEF stimulation showed an increasing trend, suggesting tighter connection between fibroblast and the surrounding microenvironment (Supplementary Fig. 4j). Birk et al. found that procollagen secreted by the cell could organize into fibrous structure and became the framework for epithelialization^40^. It suggested that these collagen fibers bundles surrounding the cells were likely to be evidence of the deposition of ECM, combined with the quantitative results of COL1A2. More collagen deposition will promote migration, thereby enhancing wound healing. Thus, μsPEF stimulation could not only influence the arrangement of collagen fibers in a 3D environment, but it can also influence the contractility of fibroblast and thus rearrange the ECM.

Based on our findings of the effects of μsPEF *in vitro*, we further explored the potential therapeutic effects of μsPEF on the wound healing of mouse skin (Fig. 4j). In the mouse wound healing model, we found that the wound area under 750 V/cm μsPEF stimulation was largely smaller than the control during the treatment (Fig. 4k, Supplementary Fig. 4k). On day 7, the wound area remained 7.16% in the treated group, significantly decreased than that of the control group (23.07%). Additionally, the μsPEF significantly inhibited the wound expansion on day 1 after stimulation, and the wound width was significant decreased than control in the following days (Fig. 4k). However, there was no significant difference in the trend of reduction in wound length (Supplementary Fig. 4l). These results indicated that the fiber alignment along the electric field lines may determine the direction of wound closure. To further characterize the wound healing process under the stimulation of μsPEF, the cross-sections of skin tissue samples were obtained and stained by hematoxylin and eosin at day 8. The result showed that epidermis of healed skin tissue was thicker in the 750 V/cm μsPEF stimulation group, indicating faster epithelial migration in the treated group, which enhanced the re-epithelialization process (Fig. 4l, m). Meanwhile, masson staining of skin tissue sections also demonstrated that the collagen fibers were well-organized in the 750 V/cm μsPEF group, consistent with *in vitro* results, and this change conformed to the features of wound healing in the remodeling stage (Fig. 4n, o). According to the above results, the application of 750 V/cm μsPEF exerted a facilitative effect on the re-epithelialization process as well as the remodeling phase. Taken together, we demonstrated that μsPEF stimulation could promote cell migration in 3D microenvironment by affecting both the cellular contractility and ECM properties, thus allowing cells to gain greater migration potential in speed and orientation, which enhanced and fastened the wound healing process (Fig. 4p).

## Conclusion

In this study, we proposed a low-frequency and short-duration μsPEF technology and explored the mechanism of μsPEF-mediated wound healing. The transient relaxation dynamics of cell electroporation is in sophisticate of usage to push the short time limit to cell response. We verified the safety of μsPEF by evaluating fibroblasts viability and temperature rise. The result presented that μsPEF had a positive effect on individual and collective cell migration, which was important for wound healing. We demonstrated that μsPEF directly regulated actin cytoskeletal reorganization and the turnover rate of focal adhesion, promoting fibroblasts migration. Furthermore, we reconstituted a three-dimensional collagen model, to more closely mimic *in vivo* microenvironment, and found that μsPEF could promote collagen fiber alignment and improve collagen gel stiffness, which provided fibroblast contact guidance and facilitated directional migration of fibroblasts. Finally, μsPEF effectively promoted the closure of the skin wound *in vivo*, and the efficiency of wound closure was higher along the electric field line, attributing to faster fibroblasts migration and the better alignment of collagen fibers. Our finding demonstrated that the precise controllable short time μsPEF stimulation can both regulate the cytoskeleton and collagen microenvironment to promote fibroblasts migration and wound healing, which was distinct from current time-consuming wound healing therapeutic methods of exogenous growth factors and biomaterials. Overall, our μsPEF technology is promising in enhancing wound healing, and the combination of this technology with different therapeutic drugs may have a better therapeutic effect. The safety and efficiency of our μsPEF device guaranteed the broad application in the field of biomaterial and regenerative medicine.

## Materials and Methods

### Cell culture

Human fetal lung fibroblast (Institute for Medical Research-90, IMR90, ATCC) cells were cultured in Minimum essential medium (MEM, 1×) (Invitrogen, USA) containing 10% fetal bovine serum (Gibco, USA), 1% glutaMAX (Gibco, USA), 1% MEM non-essential amino acids (NEAA, 100×, Gibco, USA), 1% Sodium pyruvate (100 mM, Gibco, USA) and 1% penicillin/streptomycin at 37℃ and 5% CO_2_.

### Device fabrication and electrical stimulation

In order to generate the rectangular pulsed monophasic μsPEF waveforms, solid-state Marx generator (SSMG) based on the capacitors charged in parallel and discharged in series was used^41,42^. The waveform of output pulses can be adjusted by the changed number of series capacitors^43^. Typical unipolar SSMG consisted of positive high-voltage pulses topology (Supplementary Fig. 1a). In these circuits, switches S_a1_-S_an_ and S_b1_-S_bn_ formed several half-bridge circuits, where switches S_b1_-S_bn_ controlled the charge of the energy storage capacitors and switches S_a1_-S_an_ controlled the discharge of the Marx generators. During the charging phase, switches S_b1_-S_bn_ were turned ON while S_a1_-S_an_ were OFF. All capacitors were charged in parallel through diodes D_1_-D_n_. During the dead-time, all switches were OFF, which can inhibit short-circuit of half-bridge circuits. At the discharging stage, switch S_a1_ controlled the discharge of C_1_, S_a2_ controlled the discharge of C_2_, and so on. The corresponding capacitors could be coupled in series, and a high-voltage pulse can be produced on the load. ΔU_*Load*_ was the voltage drop of the output pulses on resistive loads, which affected the selection of energy storage capacitors. In this part, the stacking of SSMGs with positive pulses topologies were manufactured, and a novel unipolar high-voltage topology was proposed.

The set-up of the PEF stimulation chamber consisted of electrodes secured to the top lid of a cell/tissue culture chamber that fitted on a standard 6-well cell/tissue-culture plate, facilitating handling and sterilization and reducing medium evaporation (Supplementary Fig. 1b). The electrodes were made from red copper, which is generally known for its excellent corrosion resistance, passivation capacity, and biocompatibility, and can also be used in promoting incisional wounds healing ^44^. To supply fibroblast with capacitively coupled alternating electrical fields, pairs of electrodes with dimensions 20 mm × 20 mm × 3 mm, separated by distance of 10 mm and 20 mm respectively for generating different electric fields intensity, were soldered vertically to the printed circuit board (PCB), fixed to the cover of the 6-well plate, and connected to a self-made pulse power supplier (six-stage solid-state Marx modulators). For sterilization, the chamber was soaked in 70% ethanol overnight, washed in sterile calcium- and magnesium-free phosphate buffer saline (PBS), and then exposed to ultraviolet (UV) light in a cell culture laminar flow hood overnight.

The pulse power supplier was programmed to generate square monophasic waves of 1500 V to the entire setup, and the waveforms were obtained from probe P6015A (Tektronix, ltd) and oscilloscope MS0-X3204 (bandwidth 200MHz, Agilent, ltd). Then square waveform electrical pulses with pulse width of 20 μs and pulses duration of 5 s were delivered into each well at a frequency of 10 Hz with variable electric intensity. The electrical stimulation setup was connected to an oscilloscope to ensure that the electrical pulses delivered to the culture wells were of the mentioned pulse duration, frequency, and voltage. All electrical stimulations were performed 24 h after cell seeding. The medium was replaced immediately after the electrical stimulation. And the control group was subjected to the same treatment without electrical stimulation.

### Cell viability and cytotoxicity

In order to identify the cell viability after μsPEF stimulation, cell viability assay kit (Beytotime) including fluorescent live-cell staining dyes (Calcein Green AM) and fluorescent dead-cell staining dyes (PI) was selectively performed before and after electrical stimulation. The cell viability was calculated based on the following equations:

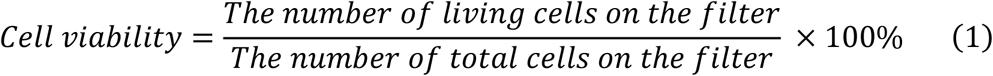

To confirm that the oxidation–reduction and electrochemical reactions of the metallic electrodes in the μsPEF exposed chamber were not cytotoxic, cell viability and metabolic activity were assessed by 3-(4,5-dimethylthiazol-2-yl)-2,5-diphenyltetrazolium bromide (MTT) assay (MTT Cell Proliferation and Cytotoxicity Assay Kit; Sangon Biotech, Shanghai, China) after cells were exposed to PEF for 1 and 3 days. Absorbance was read at 570 nm wavelength, using an Infinite 200PRO NanoQuant device, with TECAN i-control™ software (Tecan, Crailsheim, Germany). We calculated fold change in optical density of electrically stimulated groups, relative to control.

### Determination of electroporation

To visualize the electroporation on the cell membrane exposed to μsPEF the electro-permeabilization of plasma membrane was measured by cellular uptake of the nucleic acid fluorescent dyes propidium iodide (PI). The choice of fluorescent dye was based on its impermeability for live cells, and thus it can only be loaded into cells under successful electroporation. Fibroblast cells at cell density of 1×10^5^ cells mL^-1^ were cultured in 6-well plates. After 15 min incubation, the cells were stimulated in a PBS containing 12.5 μM PI (A601112, Sangon, Shanghai) under the above electrical parameters. And the fluorescence intensity changes were detected through a spinning disk confocal fluorescence microscope (W1, Nikon). The cell electroporation efficiency was calculated based on the following equations:

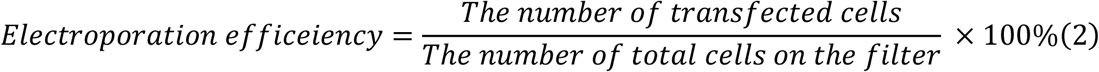

To further characterize the electroporation in cell membrane surface morphology, GeminiSEM 500 (ZEISS, German) was used by the following parameters: work distance = 4 mm, accelerating voltage = 1 kV, magnification = 20000 ×. Cells were seeded in monocrystalline silicon chip (1 cm × 1 cm) and cultured in 24-well culture dishes with medium for 2 days, followed by immediate fixation after electrical stimulation, overnight at 4℃. Dehydration was performed the next day. All samples were placed on an aluminum sample dish and coated with 2 nm gold for optimal imaging.

### Live-cell imaging

For long-term live cell imaging, IMR90 cells were plated on fibronectin (25 μg/mL)-coated glass-bottom cell culture dishes and maintained in complete culture medium without penicillin and streptomycin at 37 ℃ throughout the imaging process. For nuclei staining, Hoechst 33258 dye (#94403, Sigma, USA) was added to complete culture medium at a final concentration of 1 μg/mL, and cells were incubated at 37°C for 30 minutes before the acquisition of images. For F-actin staining by SiR-Actin Kit (#CY-SC001, Cytoskeleton, Inc.), the staining solution was prepared by diluting SiR-actin and verapamil (a broad-spectrum efflux pump inhibitor) to the desired concentration 1 μM and 10 μM respectively in cell culture medium (MEM + 10% FBS) and vortexed briefly. When cells reached the desired density, the culture medium was replaced by the staining solution and incubated at 37°C in a humidified atmosphere containing 5% CO_2_ at least 6 hours. Time-lapse imaging was recorded using an inverted fluorescence microscope (DMi8, Leica). Then the dynamic images were analyzed by the Trackmate plugin in FIJI ImageJ to obtain the cell tracking information^45,46^. Cell tracks were filtered to only keep those with good fidelity, as measured through Trackmate’s ‘quality’ filter. Mean squared displacement (MSD) is a common metric for measuring migration speed and distance traveled to characterize cell persistence^47^. The MSD is given by

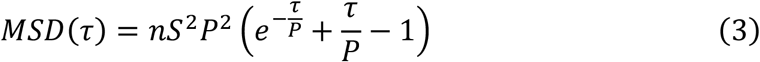

where *P* is the persistent time, *S* is the cell speed, n is the dimension of the extracellular space (which can be 1D, 2D, and 3D)^48-51^, and τ is the time lag between positions of the cell. The autocorrelation function of the cell velocity vector for the classical persistent random walk (PRW) model of cell migration exhibits a single exponential decay:

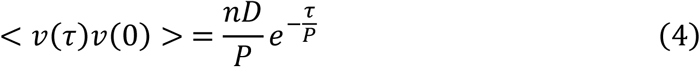

where *D* is the cell diffusivity. In the 2D, the velocity direction is described by an angle with respect to a laboratory frame, *θ*. The change in angle over a small-time interval, *dθ*, is a random variable given by a uniform distribution with a peak near *dθ* = 0. Typically, Eq. 1 was used to fit measured MSD data. The statistics of *dθ* and the time lag dependence of the velocity autocorrelation function (Eq. 2) are generally not examined in detail^52^. A customize MATLAB (MathWorks, Natick, MA) script was used to analyze the trajectories, velocities, diffusion coefficients, and migration angles. Displacement vectors of the F-actin were generated using particle imaging velocimetry PIV lab v2.54^53-55^.

### Monolayer wound healing assay

*In vitro* wound healing assays were performed as previously described ^56^. The fibroblasts were seeded (5×10^4^ cells/well) on fibronectin (10 μg/mL)-coated glass-bottom cell culture dishes. When cells reached confluence, a scratch (wound) was created on each confluent monolayer using a 200 µL sterile pipette tip perpendicular to the bottom of the dish. This generated a wound about 0.44 to 0.50 mm in width. After exposure to electric stimulation, the cultures were refreshed with culture medium and maintained at 37°C and 5% CO_2_. Images of each wound were taken by an inverted light microscope (DMi8, Leica) at 1 min interval for 24 hours following the scratch of the wound. Wound closure (cell migration) was investigated by measuring the area of the wound using the NIH ImageJ macro tool. And the PEF-exposed cultures were compared with the non-exposed ones.

### 3D Cell culture in collagen matrix

Collagen I from rat tail tendons (#354236, Corning) was used for the 3D culture. The stock solution was mixed with sterile 10× phosphate buffered saline (PBS), sterile distilled water (dH_2_O) and 1M NaOH to reach a final concentration of 1mg/mL. All the procedures were performed on ice to prevent any unwanted polymerization of the collagen gel. Cells were seeded between two layers of collagen gel to establish the sandwich-culture conditions^57^. Specifically, ice-cold neutralized collagen solution (150 µL) was added onto the 20 mm glass-bottom cell culture dish, and evenly distributed on the glass surface using the pipette tip, followed by incubation at 37°C and 5% CO_2_ for 25 min to allow collagen to polymerize and form fibrillar meshwork. Fibroblast cell suspension (100 µL) at density of 1.25 × 10^4^ cells/mL in desired medium was added onto the top of the polymerized collagen layer and then incubated at 37°C and 5% CO_2_ for 30 min, allowing the attachment of cells to the collagen matrix. After aspiration of culture medium, the second layer of collagen gel was formed by applying dropwise 150 µL hydrogel on the top of this sandwich culture and incubated for 30 min to crosslink into fibrillar collagen. Finally, the culture dishes were placed in a 37°C incubator for 24 hours to allow the formation of cell polarity and the migration of fibroblasts in the 3D collagen matrix.

### Immunofluorescence and imaging analysis

For immunofluorescence staining, cells were washed with PBS and fixed with 4% formaldehyde in PBS for 15 min at room temperature. Then cells were permeabilized and blocked simultaneously in 5% BSA (bovine serum albumin) and 0.3% Triton X-100 in PBS for 1 h. The primary antibodies were 1:100 diluted by Primary Antibody Dilution Buffer (1% BSA and 0.3% Triton X-100 in 1× PBS) and incubated at 4°C overnight. After washing with PBS for three times, the glass-bottom cell culture dishes were incubated with Alexa Fluor 488 or 555-conjugated secondary antibody for 1 h at room temperature. Alexa Fluor™ 488 phalloidin (#A12379, Thermo) was used to stain actin cytoskeleton. Then the coverslips were mounted with Antifade Mounting Medium with DAPI (#P0131-5ml, Beyotime). Detailed information of antibodies used in this study was as follows: Vinculin Recombinant Rabbit Monoclonal Antibody (#700062, Thermo), Anti-rabbit IgG (H+L), F(ab’)2 Fragment (Alexa Fluor® 488 Conjugate) (#4412S, Cell Signaling Technology). The cells were imaged with an inverted fluorescence microscope (Ti2-U, Nikon) and a spinning disk confocal fluorescence microscope (W1, Nikon) for z-stack scanning. The quantification of fluorescence intensity was processed by ImageJ and 3D reconstructed confocal z-stacks were performed by IMARIS 9.0 software (Bitplane, Swizterland).

### Focal adhesion dynamics analysis

Vinculin-GFP transfected IMR90 cells were placed into the incubation chamber (37°C) on the stage of a total internal reflection fluorescent microscope (TIRF, Nikon). Images were taken at 1 min interval for IMR90 cells and were processed for acquiring various parameters using Focal Adhesion Analysis Server (FAAS) (http://faas.bme.unc.edu)^58^. Min Adhesion Size was set to 2 pixels. Figures of FA number and size were plotted by GraphPad Prism 8.0.

### Fibrillar collagen alignment quantification

The collagen matrix preparation procedure was performed using the same method mentioned above. After PEF stimulation, the polymerized collagen gel was rinsed with PBS and fixed with 4% formaldehyde in PBS for 15 min at room temperature. For the detection of fibrillar collagen structures, collagen gel was visualized using second harmonic generation (SHG) and full-spectrum multiphoton fluorescence lifetime imaging system (Leica TCS SP8 DIVE FALCON). Z-stacks separated no more than 0.5 μm were collected. The excitation wavelength was 976 nm generated by 20 mW excitation power and the emission wavelength was from 483 to 493 nm. CT-FIRE module (MATLAB) was used to extract individual fiber and quantify fiber alignment in SHG images^59^.

### ELISA for wound healing-related cytokines

IMR90 cells were plated into 6-well plates and cultured for 24 h, followed by electrical stimulation. Then, the cells were cultured in serum-free medium for 48 h. The content of type I α collagen and basic fibroblast growth factor (FGF-2) in the supernatant were measured using commercially available Human COL1A2 (Collagen Type I Alpha 2) ELISA Kit (Elabscience, Wuhan, China) and Human bFGF/FGF2 (Basic Fibroblast Growth Factor) ELISA Kit (Elabscience, Wuhan, China) according to the manufacturer’s protocol respectively. Absorbance was read at 450 nm wavelength, using an Infinite 200PRO NanoQuant device, with TECAN i-control™ software (Tecan, Crailsheim, Germany). We calculated fold change in optical density of electrically stimulated groups, relative to control.

### *In vivo* wound healing assay

The experiment was divided into two groups: the control group and the ES group. A total of 7 ICR female mice (12 weeks, 30∼40 g) were randomly selected to receive general anesthesia by intraperitoneal injection of 2.5% tribromoethanol at a volume of 0.2 mL/10 g. The mice were fixed and then shaved with an animal electric shaver to remove the hair on their backs. Further hair removal was achieved with irritation-free depilation cream to fully expose the operative area. Alcohol (70%) was used to disinfect the surgical area, and then, a sterile towel was applied. A round wound (diameter: 5 mm) with full-thickness skin removal was made by a skin biopsy punch on the back of each mouse after anesthesia. Then the right wounds were stimulated by parallel plate electrodes with 5 mm spacing using the following parameters: 50 pulses of 20 μs, 20 Hz and 750 V/cm (applied voltage to electrode distance ratio), and the left control group remained untreated. The mice were stimulated at 24 hours intervals. All operations were performed under aseptic conditions. The mice were fed alone after operation and the wounds were photographed from day 0 to day 8 post-operation, followed by measuring the wound width and length. The progress of wound closure was evaluated by the relative size to day 0. ImageJ was used to measure and calculate the circumference of each wound boundary to obtain macroscopic wound healing data. All *in vivo* experiments were performed in accordance with the Principles of Laboratory Animal Care and approved by the Ethics Committee of Fudan University.

### Histological analysis

The skin at the wound site of each group of mice was cut and fixed. For the histological staining, the wound area was selected for analysis. Tissue sections were fixed overnight in 10% buffered formalin at room temperature. Next, the samples were transferred to 70% ethanol for an additional 48 h and then embedded in paraffin. The sections were stained with hematoxylin and eosin (H&E) and Masson’s trichrome. Images were collected using NanoZoomer (Hamamatsu Photonics, Japan). For each sample, at least 5 fields were randomly selected for analysis. All histological analysis were performed by two independent observers. MatFiber module (MATLAB) was used to automated fiber orientation analysis in Masson images^60^.

### Data distribution analysis

Since the basic data are not sufficient to characterize the data distribution, we designed 6 polynomial data based on the 3 basic data sets (speed, acceleration, direction), and obtain 9 high-dimensional data in total, according to the literature^61^, where they showed the advantages of polynomial data for data classification. Subsequently, the high-dimensional data are mapped to the low-dimensional space by t-distributed stochastic neighbor embedding (t-SNE)^62^, showing the cluster representations of similar datapoints in low-dimensional data space. To ensure similarity between high- and low-dimensional data spaces, t-SNE utilizes the gradient descent method to minimize the Kullback-Leibler (KL) divergence of the two probability distributions, formulated as follows:

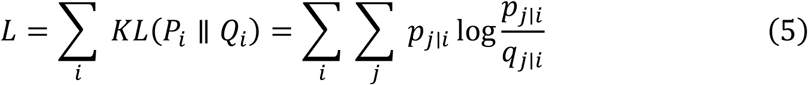

where *L* is the cost function, *KL*(·) denotes the KL calculation. The *P*_*i*_ and *Q*_*i*_ are the conditional probability distributions over all other datapoints given datapoint *x*_*i*_ and corresponding *z*_*i*_ in high- and low-dimensional spaces, respectively. Specifically, the individual conditional probabilities *P*_*j*|*i*_ and *q*_*j*|*i*_ are given by:

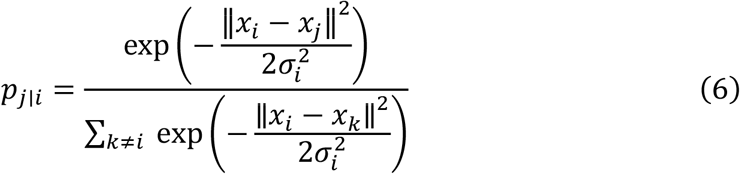

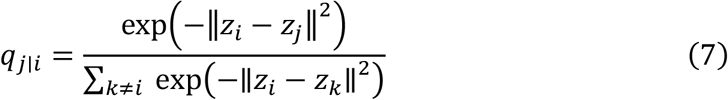

where *σ*_*i*_ is a variance of the Gaussian distribution centered on datapoint *x*_*i*_. Here, Eq. 10 is similar to Eq. 9 in which we set the variance *σ*_*i*_ to 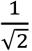. Mathematically, if the mapped points *z*_*i*_ and *z*_*j*_ correctly model the similarity between the high-dimensional datapoints *x*_*i*_ and *x*_*j*_, then the conditional probabilities*q*_*j*|*i*_ and *E*_*j*|*i*_ and will be equal.

### Quantification and statistical analysis

The number of biological and technical replicates and the number of samples were indicated in figure caption. Data were presented as mean ± standard deviation (SD) or standard error of the mean (SEM). Statistical analysis was performed by two-tailed Student’s *t*-tests between two groups or by one-way ANOVA among more groups using GraphPad Prism 8.0 (Graphpad Software Inc., USA). *p < 0.05, **p < 0.01, ***p < 0.001 and ****p < 0.0001 were considered statistically significant.

## Data availability

The authors declare that the source data supporting the findings of this study are provided with the manuscript and supplementary information files.

## Code availability

PIV lab v2.54 MATLAB code is available at https://www.mathworks.com/matlabcentral/fileexchange/27659-pivlab-particle-image-velocimetry-piv-tool-with-gui. CT-Fiber MATLAB code may be available at https://eliceirilab.org/software/ctfire/. MatFiber code may be found on GitHub at https://github.com/cardiacbiomechanicsgroup/MatFiber. All other relevant codes are available from the corresponding author on reasonable request.

## Acknowledgments

This study was supported by grants from the Pioneering Project of Academy for Engineering and Technology of Fudan University (No.gyy2018-002) and the National Natural Science Foundation of China (Nos. 31870978, 51877046, 22274026). We thank Dr. Zhanfeng Yang (Fudan University) for helping in electron microscopy measurements, Shuqing Zhang (Fudan University) for providing digital slide scanning techniques and Huiqin Li (Shanghai Jiaotong University) for providing AFM technical support.

## Author contributions

X.W.X., H.T.L., K.F.L. and Y.-J.L. conceived the idea and designed the study. X.W.X., H.T.L., Z.Y.Y., J.Q. and K.F.L. designed the stimulation device. X.W.X., W.L., Y.L.Z., Y.J.W., Y. D. and X.X.X. performed the in vitro experiments and analysis. X.W.X., Y.L.Z., Y.C.C., S.X.Y., C.H.Z., C.W. and J.T.W. performed the in vivo experiments. X.W.X., H.T.L., Z.Y.Y. and J.L., performed data analysis. X.W.X. wrote the original draft. Y.-J. Liu and K.F.L. supervised the project, wrote the manuscript and were responsible for the funding acquisition. All authors commented and reviewed the manuscript and figures.

