## Supplementray material for "Microsecond pulse electrical stimulation modulates cell migration"

**Supplementary material**

Xiao-Wei Xiang[a1], Hao-Tian Liu[a1], Wei Liu[a], Ze-Yao Yan[a], Yu-Lian Zeng[b], Ya-Jun Wang[a], Jing Liu[a], Yu-Chen Chen[a], Sai-Xi Yu[a], Cai-Hui Zhu[a], Xiao-Nan Tao[a], Chen Wang[a], Jin-Tao Wu[b], Yang Du[a], Xin-Xin Xu[a], Hai Gao[a], Yaming Jiu[c], Jiong Ma[d], Jian Qiu[a], Lingqian Chang[e], Guangyin Jing[f], Ke-Fu Liu[a*], Yan-Jun Liu[a*]

[a] Academy for engineering & technology, Shanghai Key Laboratory of Medical Epigenetics, Institutes of Biomedical Sciences, Shanghai Stomatological Hospital, and School of information science and technology, Fudan University, Shanghai, China

[b] Department of Anesthesiology, Department of Orthopedics, Ruijin Hospital, Shanghai Jiao Tong University School of Medicine, Shanghai, China

[c] The Center for Microbes, Development and Health, Key Laboratory of Molecular Virology and Immunology, Institut Pasteur of Shanghai, Chinese Academy of Sciences, Shanghai 200031, China

[d] Department of Optical Science and Engineering, Shanghai Engineering Research Center of Ultra-Precision Optical Manufacturing, Key Laboratory of Micro and Nano Photonic Structures (Ministry of Education), Green Photoelectron Platform, Fudan University, Shanghai, China

[e] Key Laboratory of Biomechanics and Mechanobiology, Ministry of Education, Beijing Advanced Innovation Center for Biomedical Engineering, School of Biological Science and Medical Engineering, Beihang University, Beijing, 100083, China

[f] School of Physics, State Key Laboratory of Photon Technology in Western China Energy, Northwest University, Xi'an, 710069, China

Corresponding authors.

### I. Supplementary information-experiments

##### A. Joule heat detection

The infrared thermal imaging in the 35 mm dish with culture medium was performed by Infrared thermal imaging camera (FOTRIC, China) before and after μsPEF stimulation at different electric field strength.

##### B. Cell adhesion and spreading assay

For the cell adhesion and spreading assay, fibroblast cells were seeded on fibronectin (25 μg/mL)-coated glass-bottom cell culture dishes. After μsPEF stimulation, each sample was rinsed with PBS, and then fixed with 4% paraformaldehyde in PBS at room temperature for 10 minutes. Afterwards each sample was stained with 40,60-diamidino-2-phenylindole (DAPI) (Beyotime) and recorded by using laser scanning confocal microscope (W1, Nikon).

##### C. Collagen gels stiffness quantification

Atomic Force Microscope (AFM) was used to detect the stiffness of the collagen gel in the control and PEF-exposed groups. This technique is a unique tool that can be operated in several different modes under different environmental conditions (e.g. air and liquid)[1]. Refering to the previous operation method[2], samples were tested using ball probe and tapping mode with Dimension FastScan AFM (Bruker, US). Force–Indentation Curves and data processing were using NanoScope Analysis 1.8.

##### D. Contractility assay

The effect of ES on the capacity of human fibroblasts to contract collagen gel was determined using a reconstituted type I collagen assay, as previously described [3]. IMR90 cells were harvested and resuspended in the desired medium at $1.5\times10^5$ cells/mL followed by mixing with 1 mg/mL collagen solution (#354236, Corning) and 1M NaOH in a tube on the ice. The mixture of cells and collagen was then seeded into a 24-well plate and cultured at 37°C with 5% $CO_2$ for 30 min. The collagen gels were detached from the well by a pipette and then the medium of appropriate volume was added into the wells. Photos of the collagen gels were taken at 0 h, 6 h and 24 h, and Fiji was used to calculate the area of collagen gels at each time point.

#### II. Supplementary information-model

##### A. Electrical equivalent circuit model (EEC)

Cell and nuclear membranes are composed of lipids, proteins, sugars, etc. The membranes are permeable to ions and thus, they have some electrical conductivity. However, membrane conductivity is much smaller, compared with the conductivity of the cytoplasm, which is dominated by aqueous electrolyte solutions. Therefore, the individual cell can also be approximated as a conductive cell membrane or nucleoplasm surrounded by a dielectric membrane. Hodgkin proposed the EEC of cell membrane by combining the properties of cell membrane and ion channels in it[61]. In the 1980s, H. Kanai et al[31] described the extracellular fluid, cell membrane, and intracellular fluid by parallel circuits of resistance and capacitance, and then proposed the equivalent circuit model of individual cells in biological tissues. And an equivalent circuit model for multiple cells in biological tissues was proposed, with $R_e$, $R_m$, $R_i$, $C_e$, $C_m$, and $C_i$ as the resistance and parallel capacitance of the extracellular fluid, cell membrane, and intracellular fluid, respectively. In the low frequency range (below 1 MHz), $R_m$ is large and can be regarded as an open circuit, while $C_e$ and $C_i$ are small and can be regarded as open circuits, which leads to the simplified equivalent circuit model (Supplementary Fig. 3e). In general, as current flows through the cell from left to right, a portion of that current will cross the cell membrane and enter the intracellular fluid. The rest of the current will flow through the extracellular fluid (Supplementary Fig. 3f). These two current paths are implemented in the EEC by using two parallel branches, for which the upper branch (capacitor $C_m$ in series with resistor $R_i$) simulates the current path through the cell membrane and the intracellular space and the lower branch (resistor $R_e$) simulates the current path through the extracellular space (Supplementary Fig. 3e). Thus, the EEC of collective cells is an array of interconnected EECs (each EEC corresponds to a single cell) which provides a modified Fricke and Morse EEC [62]. The capacitor $C_M$ is replaced by a fractional capacitor $C_{m,\ tissue}$ which is considered as the space distribution of the electrical tissue properties [63]. The simulation had shown the electric field line distribution and the field intensity distribution of single cells and collective cells at 750 V/cm and 1500 V/cm respectively (Supplementary Fig. 3g).

#### B. Numerical simulations

The numerical simulation used to qualitatively verify the electric potential distributions on individual cells and monolayers cells subjected to the same uniform electric field was performed in COMSOL 5.4 Multiphysics (Stockholm, Sweden). A simple two-dimensional geometry, consisting of three 30 μm diameter circles placed in the center, was used to model the simplified EEC. To differentiate cells in single and cells in tightly connected monolayers, the spacing between single cells was set as 40 μm, and the spacing between cells in monolayers was 29 μm. Electric potential $\phi$ can be solved according to the following equation:

$$\nabla \cdot \left( \left( \sigma + \varepsilon \frac{\partial}{\partial t} \right) \nabla \phi(x, y, t) \right) = 0 \tag{1}$$

where $\sigma$ is conductivity, and $\varepsilon$ is the relative permittivity. This equation describes the electric potential $\phi(x, y, t)$ as a function of the distribution of space and time. The conductivity of the suspension is 0.2 $S/m$, the conductivity of the cell is 0.3 $S/m$, the relative permittivity of the suspension is 80, and the relative permittivity of the cell is 154.4. These parameters designed in the model were based on previous studies[61,62].

We simplified the equation for the steady state ($\frac{\partial}{\partial t} = 0$) and uniform conductivity ($\nabla \cdot \sigma = 0$), and then we obtained the Laplace's equation

$$\nabla \cdot \nabla \phi(x, y) = 0. \tag{2}$$

The electric field intensity is the gradient of the electric potential, so

$$E = -\nabla \phi \tag{3}$$

The left wall (x=0) was provided with PEF voltage while the right wall was grounded, and the top and bottom borders were defined as an insulating boundary.

#### III. Supplementary movies

**Supplementary movie 1** Fibroblasts migration after electrical stimulation on fibronectin coated surface. Phase contrast images. Fluorescence intensity of nucleus. Scale bar 50 μm.

**Supplementary movie 2** Actin rearrangement in fibroblast without/with electrical stimulation, recorded by TIRF microscopy of SiR-actin (actin filaments) in fibroblast cells. Scale bar 20 μm.

**Supplementary movie 3** Focal adhesions turn over in fibroblasts without/with electrical stimulation, recorded by TIRF microscopy of Vinculin-GFP (focal adhesion) in fibroblast cells. Scale bar 20 μm.

**Supplementary movie 4** Collective fibroblasts migration after electrical stimulation on fibronectin coated surface. Phase contrast images. Scale bar 200 μm.

#### IV. Supplementary Figures

***Fig 1-SI The design and fabrication of the SSMG device and parameter optimization***

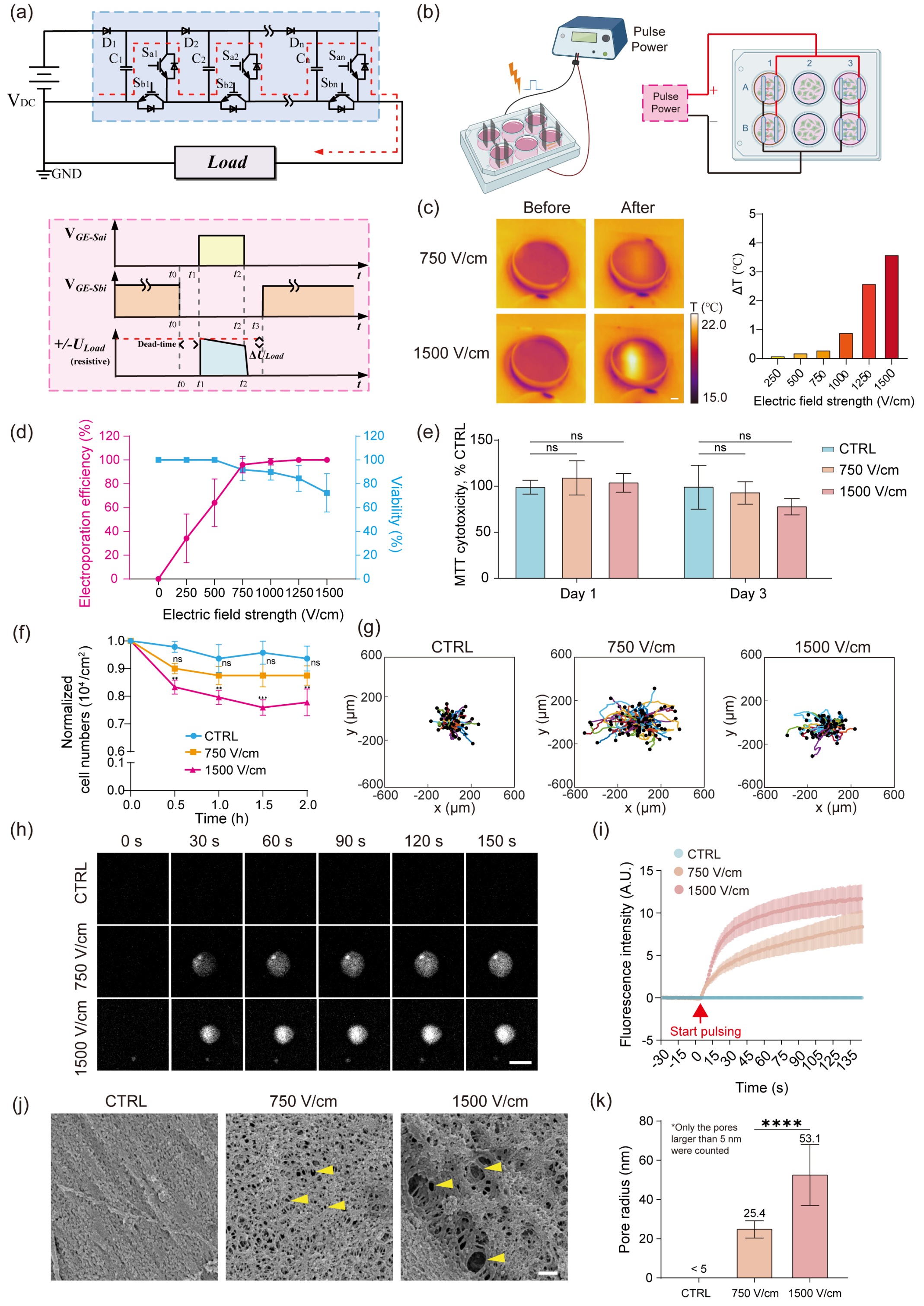


**Supplementary Figure. 1 The design and fabrication of the SSMG device and parameter optimization**

(a) Schematic diagram of n-stage solid-state Marx modulators. Top: high-voltage pulses topology. Bottom: the diagram of control signals and output pulses.

(b) Experimental diagram: SSMG was applied to two of the three columns of the 6-well plate, and the remaining column was used as control. Four pairs of electrodes were secured to the lid of a 6-well plate, and the electrodes were in contact with the bottom of the cell culture plate and fully covered by culture medium. Column 1 and 3 of the 6-well plate could generate different field strengths by varying the electrode spacing, and the field strength generated in column1 was double that in column3 when the same voltage was applied. And column 2 was used as control. Rows 1 and 2 were used as duplication. Created with BioRender.com.

(c) Infrared thermal imaging of the medium before and after the treatment with 750 V/cm and 1500 V/cm μsPEF. Bar chart showing temperature rise of the medium affected by different electric field strengths (250, 500, 750, 1000, 1250, 1500 V/cm). Scale bar, 5 mm.

(d) The electroporation efficiency and viability under different electric field strengths (n=3 for every trial).

(e) MTT assay showing the cytotoxicity of IMR90 cells after μsPEF, where exposed group had no significant difference at days 1, 3 compared with the control. Results are presented as mean ± standard deviation with 95% CI (n=5); ns>0.05 versus control.

(f) Cell adhesion ability assay showing IMR90 adhesion ability after μsPEF within 2 h by counting the stained nuclei with Hoechst 33258. Results are presented as mean ± standard error of the mean (n=5); $^{**}p<0.01$, $^{***}p<0.001$, ns>0.05 versus control by two-way *ANOVA* for multiple comparisons.

(g) Representative trajectories of cell migration on fibronectin-coated surface under different intensity μsPEF (i.e., 750 and 1500 V/cm) and control. Each plot has 100 cells trajectories, and each colored line represents the path of an individual cell in 24 h.

(h) Representative time-lapse images showing the electroporated IMR90 cells in the presence of PI at 750 V/cm and 1500 V/cm. Scale bar, 20 μm.

(i) Fluorescence intensity curves of control, 750V /cm and 1500 V/cm exposed cells ($n_{CTRL}$=8 cells, $n_{750\ v/cm}$=8 cells, $n_{1500\ v/cm}$=8 cells, error bar: mean with 95% CI).

(j) Representative SEM images of electroporated IMR90 cells exposed to750 V/cm and 1500 V/cm μsPEF. The pores left after μsPEF stimulation are indicated by yellow arrows. Scale bar, 200 nm.

(k) Bar plot showing the pore radius after electrical exposure. The inherent pores of the cell are not counted as electroporation induced pores, i.e., pores below 5 nm. Results are expressed as mean ± standard deviation with 95% CI (n=20). Statistically significant results are shown as $^{****}p<0.0001$.

*Fig 2-SI μsPEF enhanced migration regulated by cytoskeleton rearrangement and focal adhesion turnover rate*

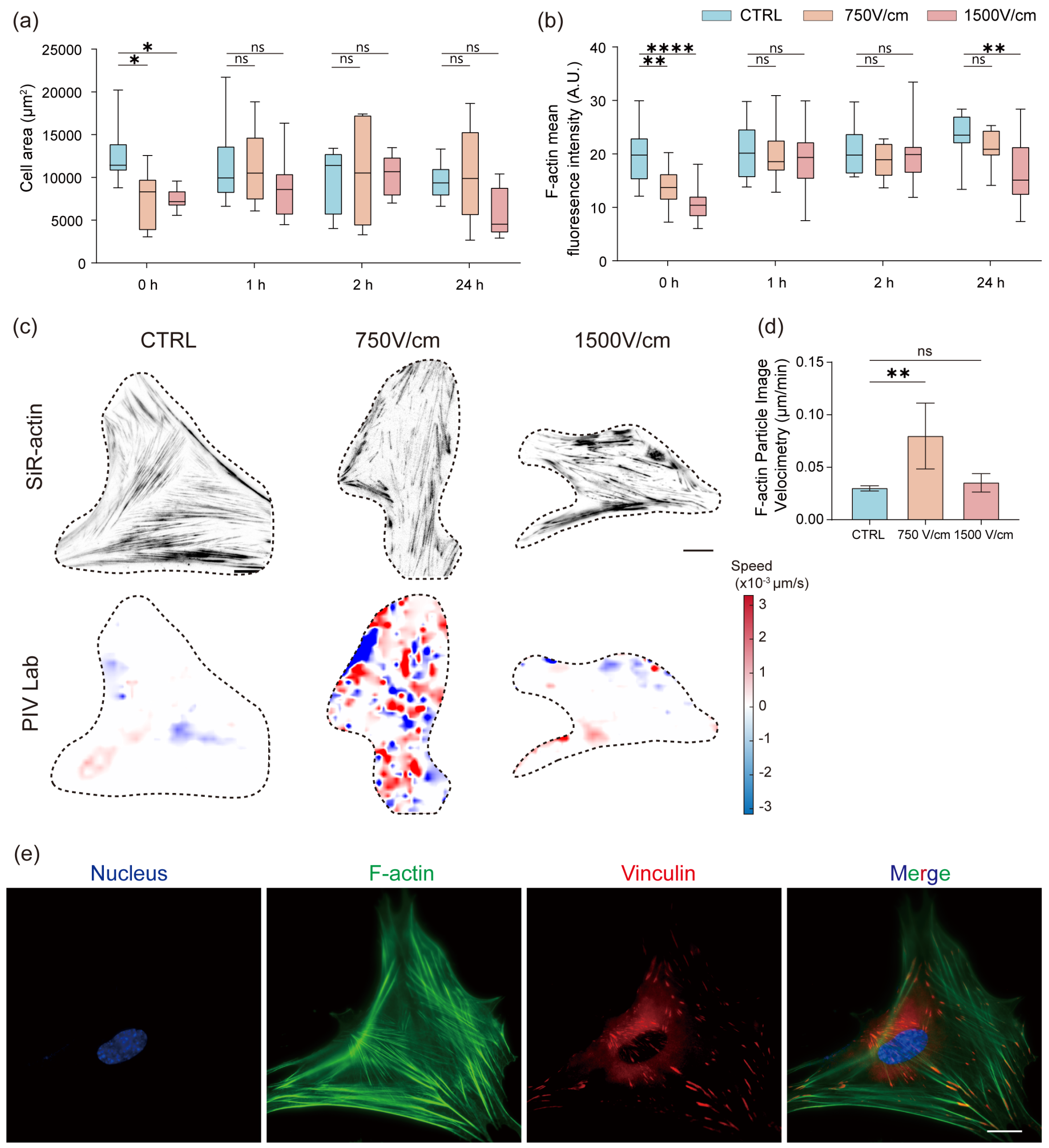


**Supplementary Figure. 2 μsPEF enhanced migration regulated by actin cytoskeleton rearrangement and focal adhesion turnover rate**

(a) Influence of the cell spreading area after μsPEF (i.e., 750 and 1500 V/cm) stimulation at different time points (i.e., 0, 1, 2 and 24 h). Results are presented as mean ± SEM (n=8); $^{*}p<0.05$, $ns_{750\ V/cm}=0.9974$ and $ns_{1500\ V/cm}=0.2628$ at 1h, $ns_{750\ V/cm}=0.9348$ and $ns_{1500\ V/cm}=0.9995$ at 2h, $ns_{750\ V/cm}=0.9273$ and $ns_{1500\ V/cm}=0.0570$ at 24h versus control by two-way *ANOVA* for multiple comparisons.

(b) Influence of the cell cytoskeleton after μsPEF (i.e., 750 and 1500 V/cm) stimulation at different time points (i.e., 0, 1, 2 and 24 h). Results are presented as mean ± SEM with 95% CI (n=8); $^{*}p<0.05$, $^{**}p<0.01$, $ns_{750\ V/cm}=0.9987$ and $ns_{1500\ V/cm}=0.6853$ at 1h, $ns_{750\ V/cm}=0.5774$ and $ns_{1500\ V/cm}=0.8278$ at 2h, $ns_{750\ V/cm}=0.5142$ at 24h versus control by two-way *ANOVA* for multiple comparisons.

(c) Top: Representative fluorescence images of IMR90 cells express F-actin upon μsPEF exposure (i.e., 750 and 1500 V/cm). Bottom: Representative images of F-actin particle image velocimetry analyzed by PIV Lab, where the direction vector of movement of each particle is distinguished by blue and red, and the speed magnitude of movement is distinguished by the shade of blue and red. Scale bar, 20 μm.

(d) Bar charts showing F-actin particle image velocimetry of control and cells under μsPEF. Results are expressed as mean ± SEM with 95% CI (n=4 cells). $^{**}p<0.01$, ns=0.8944 versus control by one-way *ANOVA* for multiple comparisons.

(e) Representative fluorescence images of untreated IMR90. Scale bar, 20 μm.

*Fig 3-SI Collective cell directional migration in response to μsPEF*

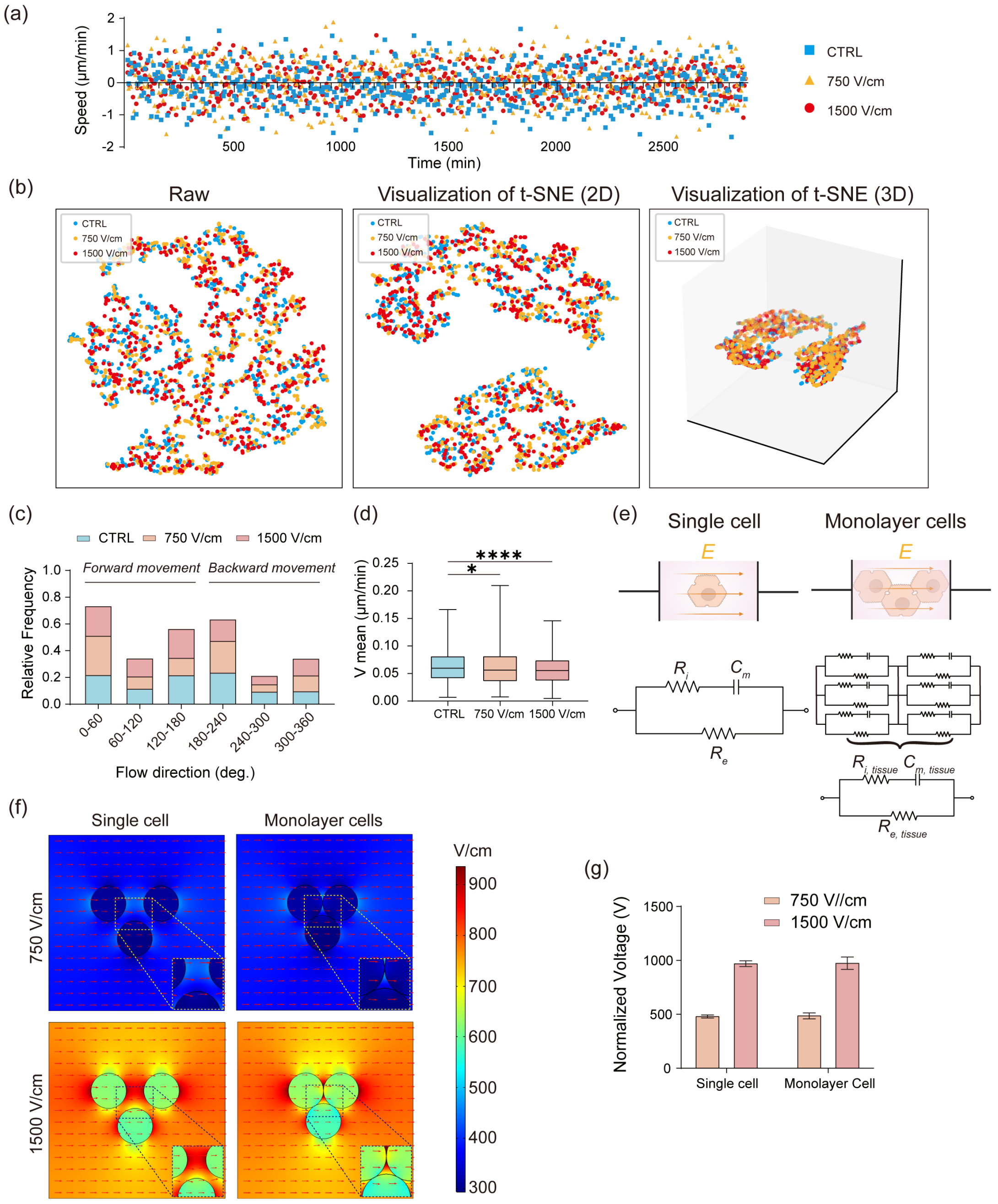


**Supplementary Fig. 3 Collective cell directional migration in response to μsPEF**

(a) The graph shows the time course of the migration speed for monolayer cell (■, ▲, ▼, represent control group, 750 V/cm treated group and 1500 V/cm treated group)

(b) t-SNE visualizations on the dataset of collection migration after μsPEF exposure (i.e., 750 and 1500 V/cm). Left: Original input space; Middle: After adding the migration flow direction factor; Right: 3D visualization.

(c) Flow direction frequency distribution of collective cell migration after μsPEF exposure (i.e., 750 and 1500 V/cm). Angle in degrees, 0 = right, 90 = up, 180 = left, 270 = down, 360 = right.

(d) Filter the mean velocity distribution of forward movement (0°≤ flow direction angle ≤180°) in control group, 750 V/cm group and 1500 V/cm group. Results are presented as mean ± SEM with 95% CI (n=3). $^{*}p<0.05$, $^{****}p<0.0001$ by one-way *ANOVA* for multiple comparisons.

(e) Cell, tissue, and electrical equivalent circuit (EEC). Top left: single cell. Bottom left: EEC of the cell. Top right: monolayer cell of biological tissue. Bottom right: EEC of the monolayer cell.

(f) Numerical simulation of the electric field distribution and direction between single cell and monolayer cell under μsPEF exposure (i.e., 750 and 1500 V/cm). Color spectrum indicates electric field intensity, with areas of low intensity in blue and high intensity in red.

(g) Bar charts showing internal electric field strength distribution of single cell and monolayer cell under μsPEF exposure (i.e., 750 and 1500 V/cm). Results are expressed as mean ± SEM with 95% CI (n=3 experiments). $ns_{750V/cm\_single\ cell\ versus\ 750V/cm\_monolayer\ cell}=0.9753$, $ns_{1500V/cm\_single\ cell\ versus\ 1500V/cm\_monolayer\ cell}=0.9857$ by unpaired Student's *t* test.

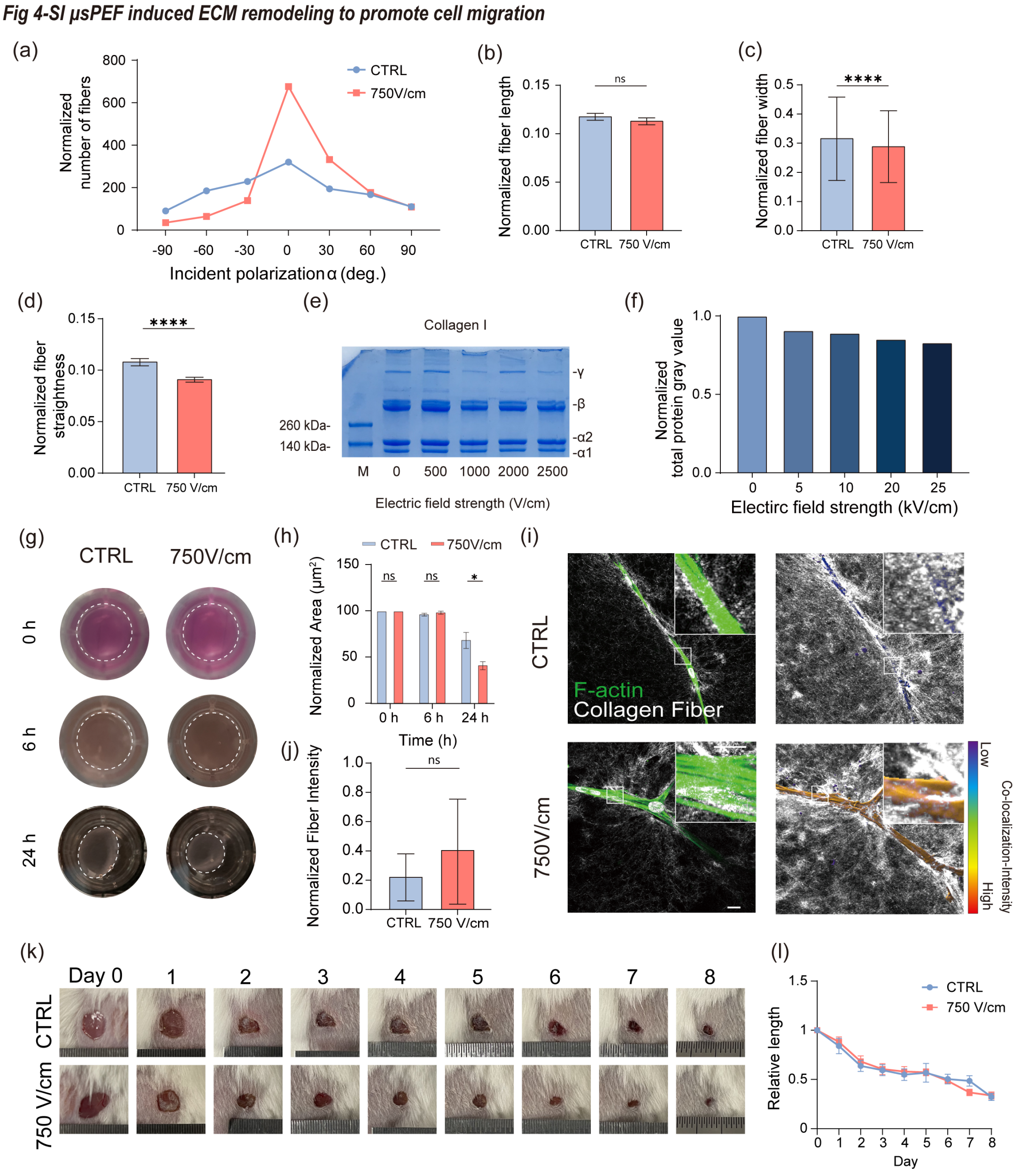


**Supplementary Fig. 4 μsPEF induced ECM remodeling to promote cell migration**

(a) Line chart showing the orientation of collagen fibers with or without 750 V/cm μsPEF treatment.

(b)~(d) Bar chart showing quantitative analysis of collagen fiber distribution with various parameters including fiber length, width and straightness. Results are presented as mean ± SEM with 95% CI ($n_{length\_CTRL}$=1058 fibers, $n_{length\_750\ V/cm}$=1051 fibers; $n_{width\_CTRL}$=990 fibers, $n_{width\_750\ V/cm}$=1051 fibers, $n_{width\_1500\ V/cm}$=2277 fibers, $n_{straightness\_CTRL}$=990 fibers, $n_{straightness\_750\ V/cm}$=1228 fibers,). $^{***}p<0.0001$, $ns_{750V/cm}$=0.7720 versus control by unpaired Student's *t* test.

(e)~(f) Comparison of collagen composition and total concentration change under different intensity μsPEF (i.e., 500, 1000, 2000 and 2500 V/cm) treatment. SDS-PAGE pattern of collagen gel on 12.5% gel (e), M for protein marker. Bar chart showing quantitative analysis of total collagen concentration.

(g)~(h) IMR90 cells are cultured in 3D collagen gels with or without μsPEF treatment. The morphology of cell-collagen gel mixture is recorded at 0, 6, and 24 h. Scale bar, 5 mm. Collagen gel contraction is quantified by the measurement of the gel area (n=3 experiments).

(i) Immunofluorescence analysis of IMR90 and collagen fiber distribution in 3D matrix upon applied different intensity µsPEF after 24h. Left: Representative fluorescence images of IMR90 in 3D collagen matrix with applied electrical stimulation. Right: Quantification of the intensity of colocalization of F-actin and collagen fiber by IMARIS. Color spectrum indicates intensity of colocalization, with areas of low intensity in blue and high intensity in red. Scale bars, 30 µm and 15µm (zoom).

(j) Bar chart showing the intensity of colocalization of fiber and cell under different intensity µsPEF after 24h. Results are expressed as mean ± standard error of the mean with 95% CI (n=3). $ns_{750\ V/cm}$ = 0.7268 versus control by unpaired Student's *t* test.

(k) Representative chronological wound images of each group are shown up to 8 d.

(l) Changes of wound length. Lines and error bars indicate mean and standard error of mean (n=7).

1 Stylianou, A. & Stylianopoulos, T. J. B. Atomic force microscopy probing of cancer cells and tumor microenvironment components. **6**, 33-46 (2016).
2 Stylianou, A., Gkretsi, V. & Stylianopoulos, T. J. M. Atomic force microscopy nano-characterization of 3D collagen gels with tunable stiffness. **5**, 503-513 (2018).
3 Semlali, A., Chakir, J., Rouabhia, M. J. J. o. T. & Environmental Health, P. A. Effects of whole cigarette smoke on human gingival fibroblast adhesion, growth, and migration. **74**, 848-862 (2011).
